# Systematic comparison of dCas9-based DNA methylation epimodifiers over time indicates efficient on-target and widespread off-target effects

**DOI:** 10.1101/2025.03.15.641804

**Authors:** Majid Pahlevan Kakhki, Fatemeh Rangani, Ewoud Ewing, Chiara Starvaggi Cucuzza, Galina Zheleznyakova, Maria Kalomoiri, Tejaswi Venkata S. Badam, Ruxandra Covacu, Ioanna Andreou, Maria Needhamsen, Lara Kular, Maja Jagodic

## Abstract

CRISPR/dCas9-based epigenome editing systems, including DNA methylation epimodifiers, have greatly advanced molecular functional studies revolutionizing their precision and applicability. Despite their promise, challenges such as the magnitude and stability of the on-target editing and unwanted off-target effects underscore the need for improved tool characterization and design. We systematically compared specific targeting of the *BACH2* gene promoter and genome-wide off-target effects of available and novel dCas9-based DNA methylation editing tools over time. We demonstrate that multimerization of the catalytic domain of DNA methyltransferase 3A enhances editing potency but also induces widespread, early methylation deposition at low-to-medium methylated promoter-related regions with specific gRNAs and, interestingly, also with non-targeting gRNAs. A small fraction of the methylation changes associated with transcriptional dysregulation and mapped predominantly to bivalent chromatin associating both with transcriptional repression and activation. Additionally, specific non-targeting control gRNA caused pervasive and long-lasting methylation-independent transcriptional alterations particularly in genes linked to RNA and energy metabolism. CRISPRoff emerged as the most efficient tool for stable targeting of the *BACH2* promoter, with fewer and less stable off-target effects compared to other epimodifiers but with persistent transcriptome alterations. Our findings highlight the delicate balance between potency and specificity of epigenome editing and provide critical insights into the design and application of future tools to improve their precision and minimize unintended consequences.

## Introduction

Genome editing systems have been extensively employed for functional studies that were revolutionized with the emergence of CRISPR/Cas-based techniques^1^. Given the risks associated with conventional CRISPR/Cas systems, the deactivated Cas (dCas) offered possibilities to target the genome by modulating its regulation through alterations of epigenetic states without DNA disruption^2^. Moreover, dCas-based systems effectively addressed the challenge of targeted manipulation of epigenetic modifications during transcriptional regulation due to the ease of design and site-specific targeting. The design involves dCas fused to an epigenetic modifier protein domain (dCas-epimodifier) that is guided by a short locus sequence-specific guide RNA (gRNA) to the genomic locus where it modulates epigenetic marks^3^.

The development of early assays to quantify DNA methylation, combined with advancements in high-throughput technologies, has enabled robust and precise genome-wide methylation profiling even in low-input and archival samples. These innovations have made DNA methylation one of the most extensively studied epigenetic marks in mammals^4,5^. DNA methylation at regulatory regions typically associates with gene repression^6^ which is achieved by different mechanism including interfering with the binding of sequence-specific and methylation-sensitive transcription factors (TFs) to promoters, enhancers or insulators^7,8^. On the contrary, gene body methylation positively correlates with gene expression^9^ and has been shown to play a role in transcriptional elongation and co-transcriptional splicing^10–12^. Nevertheless, the context dependent impact of DNA methylation on gene expression is still far from being well understood. It is therefore not surprising that functional consequences of methylation changes that are associated with human diseases remain largely unknown. To rigorously determine the impact of DNA methylation, it is necessary to study transcriptional and cellular consequences of deposition or removal of methylation marks. Traditional methods for perturbing methylation levels involve non-specific genetic or pharmacological tools with pleiotropic effects^13^, while the development of CRISPR-based tools, owing to its flexibility, specificity and efficiency, opened possibilities for desired manipulation of methylation levels in a specific locus^14^.

A simple fusion of dCas9 with DNA (cytosine-5)-methyltransferase 3A (DNMT3A) has been demonstrated to efficiently increase the methylation at a desired locus leading to decreased expression of the targeted gene *in vitro*^3^. Moreover, fusion of dCas9 with DNMT3A or ten-eleven translocation methyl cytosine dioxygenase 1 (TET1) has been shown to effectively remodel locus-specific methylation also *in vivo*^15^. Adding DNA (cytosine-5)-methyltransferase 3-like (DNMT3L) further enhances the *de novo* methylation by DNMT3A, although it also leads to unspecific changes in the predicted off-target binding sites^16^. Non-specific methylation of the human genome was observed also with bacterial DNMT although the mutated version called MQ1^Q147L^ revealed a fast and more precise targeting of DNA methylation^17^. Nevertheless, *de novo* off-target methylation with DNMTs can be widespread^18^. Increased genome-wide on-target/off-target effects ratio has been achieved using the SunTag system that enables local amplification of the epimodifier^19^. Interestingly, a more recent design called CRISPRoff, showed an efficient and stable methylation with a capability of silencing the genes lacking canonical CpG islands^20^. This design combines DNMT3A and DNMT3L with the human Kruppel-associated box (KRAB) repressor domain, which recruits a set of epigenetic silencers including histone methylases in a single fusion protein. Recently, a multi gRNA approach was utilized to remodel methylation and multiple genomic loci simultaneously^21^.

In general, the epimodifier system designs are aiming at maximizing the on-target effect with minimal off-target effects. However, due to variations in the epimodifier constructs, delivery approaches, methodologies for DNA methylation assessment, different strategies for off-target evaluation, target locations and cellular systems, it is challenging to compare available systems. Furthermore, the sensitive nature of DNA methylation, combined with the activity of endogenous *de novo* DNMTs, pose significant challenges to accurately assess exogenous dCas9-methyltransferase activity within the nucleus^18^.

The aim of this study was to provide a comprehensive systematic comparison of the on-target/off-target effects across several available and novel epimodifiers over time. Moreover, we investigated the potential of a strategy based on multimerization of epimodifier domains.

## Results

### Efficiency and specificity of DNA methylation editing at predicted gRNA-binding loci

To compare all the available dCas9-based DNA methylation editing constructs (hereafter called epimodifiers), we subcloned the catalytic domain of DNMT3A (dCas9-3A), M.SssI (dCas9-M.SssI), SunTag (dCas9-SunTag) as well as deactivated DNMT3A (dCas9-d3A) sequences into the same plasmid backbone composed of the catalytically inactive S. pyogenes Cas9 (dCas9) (**Fig. 1a**). To investigate the potential of a strategy based on multimerization of epimodifier domains, we generated additional constructs with dCas9-DNMT3A fused to DNMT3A (dCas9-3A3A), DNMT3L (dCas9-3A3L) or KRAB (dCas9-3A-KRAB). Therefore, all epimodifiers incorporated the same plasmid backbone expressing GFP and gRNA under the CMV and U6 promoter, respectively, while only the methylating module varied (**Fig. 1a**). We verified the expression of dCas9 fusion proteins by Western blot (**Suppl. Fig. 1**). In addition to these constructs, we used the recently reported CRISPRoff construct harboring DNMT3A-DNMT3L-dCas9-BFP-KRAB (CRISPRoff) without changing the original backbone^20^.

**Figure 1.**
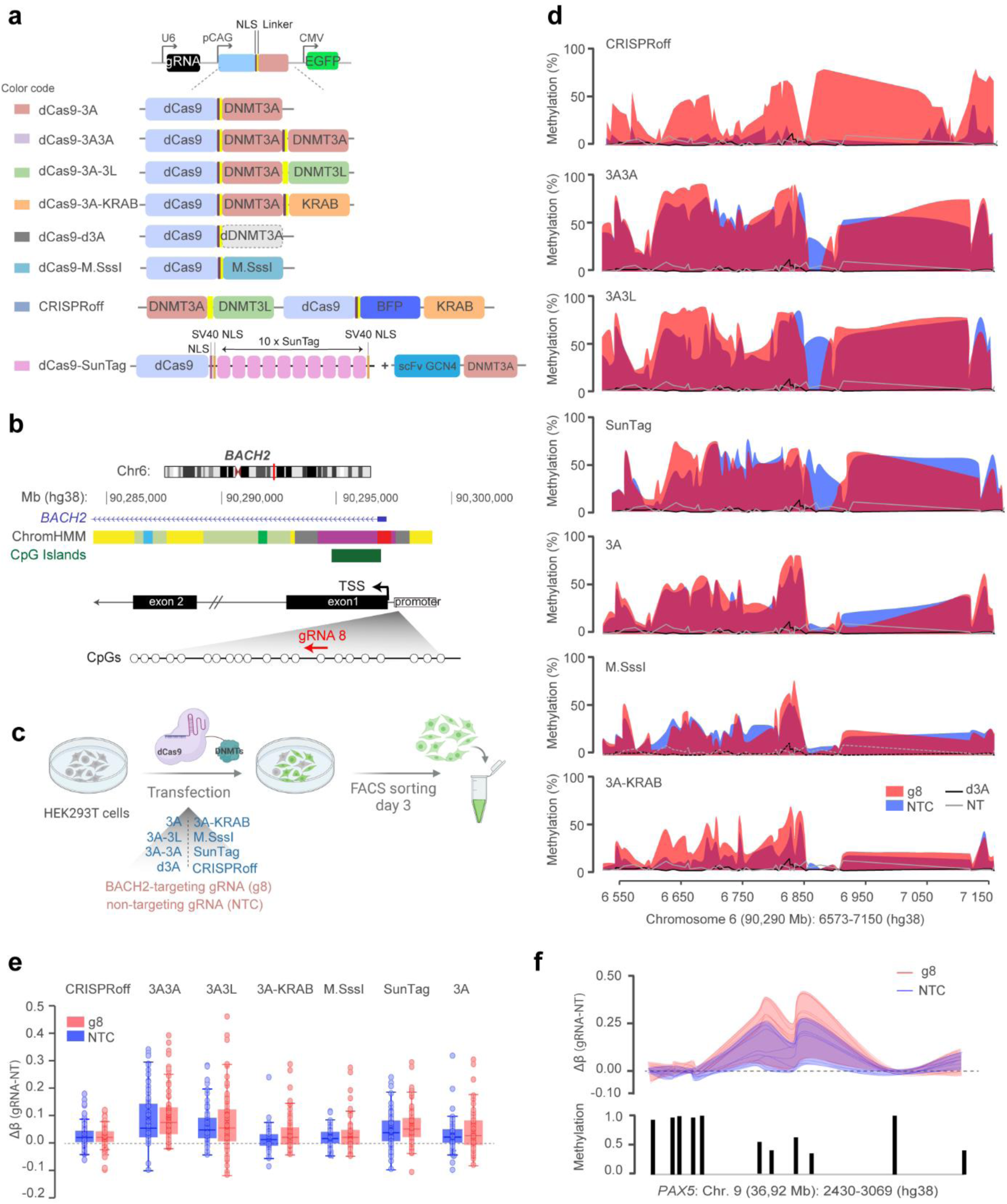
Early methylation editing efficiency of the dCas9-epimodifiers. **a.** Schematic of the design of the dCas9-epimodifiers. **b.** Genomic localization of *BACH2* promoter targeted by the specific *BACH2*-targeting gRNA 8 (g8). **c.** Experimental design where HEK293T cells were transfected with plasmids encoding gRNA-dCas9-epimodifier-EGFP (3A, 3A3L, 3A3A, 3A-KRAB, mut3A, M.SssI, SunTag) or CRISPRoff (co-transfected with a gRNA-encoding plasmid). Successfully transfected cells were sorted 3 days post transfection (p.t.) by GFP-based fluorescent activated cell sorting (FACS). **d.** DNA methylation of CpGs in *BACH2* promoter 3 days p.t. with epimodifiers expressing either *BACH2*-targeting gRNA8 (g8) or non-targeting control gRNA (NTC). Methylation levels in the control cells including cells expressing dCas9-deactivated DNMT3A (d3A) or non-transfected cells (NT) are depicted by the black and grey lines, respectively. **e.** Tukey plot, comparison Δβ (gRNA-NT) of the methylation changes induced by the g8 versus NTC across all tools. Boxplots summarize the distribution, with positive Δβ indicating increased methylation. **f**. Example of methylation changes at one of the predicted off-target loci (*PAX5*: Chr. 9 36,92,2430-36,92,3069, hg38), with methylation change induced by g8 or NTC compared to NT control (upper graph). The basal methylation level at each CpG is represented by the histogram plot (lower graph). Methylation data, assessed by targeted methylation sequencing, are available in Supplementary Table 3.

To test the efficiency of DNA methylation deposition, we transfected HEK293T cells with each construct containing the previously validated gRNAs targeting either the *BACH2* (g8)^3^ (**Fig. 1b**) or *CXCR4* (g3)^16^ promoter and a non-targeting control gRNA (NTC) (**Suppl. Table 1**). To address both on-target and potential off-target effects, we designed targeted methylation-sequencing (TMS) assays that cover 84 and 64 CpGs at the *BACH2* (586 bp) and *CXCR4* (750 bp) locus, respectively, as well as CpGs in loci predicted as potential off-target regions throughout the genome based on homology with gRNA sequence (see Methods) (**Suppl. Table 2**). Methylation was assessed in transfected GFP-expressing cells sorted three days post-transfection (p.t.) (**Fig. 1c**, **Suppl. Fig. 2a**). We observed a marked increase in DNA methylation after delivery of each epimodifier carrying *BACH2*-targeting g8 gRNA, although with varying efficiency (**Fig. 1d**). Starting from 0-13% basal methylation (β-values) at the *BACH2* promoter CpGs, dCas9-3A3A and dCas9-3A3L epimodifiers induced the most pronounced DNA methylation (average β = 68%) followed by CRISPRoff, dCas9-SunTag and dCas9-3A (average β = 40%), while dCas9-M.SssI and dCas9-3A-KRAB exerted more moderate effects (average β = 26%) (**Fig. 1d**, **Suppl. Table 3)**. Expectedly, dCas9-d3A containing deactivated DNMT3A had no impact on DNA methylation and resembled the basal unmethylated pattern of non-transfected cells. Notably, the scrambled NTC gRNA used as a control of the editing specificity exhibited similar effects to the majority of g8-containing constructs apart from CRISPRoff that presented with low NTC-driven methylation (**Fig. 1d**, **Suppl. Table 3**). The potent effect of both g8 and NTC gRNAs on the *BACH2* promoter could be validated by pyrosequencing of 13 overlapping CpGs (**Suppl. Fig 2b**). Investigation of the loci outside *BACH2* predicted to potentially be targeted based on the gRNA sequence homology, demonstrated sizeable (up to 46% changes vs. d3A) methylation deposition exerted by both g8 and NTC gRNAs (**Fig. 1e**). Pronounced off-target methylation deposition was, as expected, more likely to occur in regions of the genome that displayed low to medium basal methylation levels (**Suppl. Table 3**), as exemplified for a locus in the *PAX5* intronic sequence (**Fig. 1f**). Of note, the off-target effects were not sequence-specific and occurred also at loci not predicted to be targeted based on homology with gRNA sequence. Accordingly, analysis at an independent locus in the *IL6ST* gene promoter confirmed a significant methylation deposition by both dCas9-3A and dCas9-3A3A with g8 (*BACH2*), g3 (*CXCR4*) and NTC gRNAs (**Suppl. Fig. 2b**). Similar unwanted on- and off-target effects could be observed at the *CXCR4* locus using *CXCR4*-targeting (g3) or non-targeting gRNAs in both pyrosequencing and TMS data (**Suppl. Fig. 2b-d, Suppl. Table 4**).

These findings indicate that multimerization of epimodifier domains confers a more potent editing ability compared to monomeric units and further revealed a strong pattern of unwanted on-target and off-target DNA methylation deposition early after transfection, particularly at regions with low basal methylation.

### Impact of gRNA-dCas9-DNMT3A complex on methylation editing specificity and efficiency

The undesirable effect of the NTC gRNA mirrored the editing efficiency of the respective targeting gRNA construct, suggesting that the unspecific effects depend on the presence of the enzymatic domain. It has been shown that formation of the dCas9-gRNA ribonucleoprotein (RNP) complex is a crucial step before the genome searching process can begin^22^. To investigate the impact of the gRNA/dCas9 complex formation on gRNA-independent methylation effects, we utilized expression cassettes containing the gRNA scaffold without inserting a specific gRNA sequence (scaffold cassette, lacking a PAM site, **Fig. 2a**). Additionally, we designed a new cassette by completely removing the U6-gRNA-scaffold sequence (empty cassette, **Fig. 2a**) that we used to transfect cells (**Fig. 2b**). Indeed, the methylating ability of dCas9-3A was conditioned on the formation of a gRNA-dCas9 complex, since removing the gRNA scaffold impeded unwanted off-target methylation deposition (**Fig. 2c**). Moreover, the methylation potency correlated with the levels of available epimodifier, with the highest deposition in GFP+++ and lowest in GFP+ sorted fraction (**Fig. 2c,d**).

**Figure 2.**
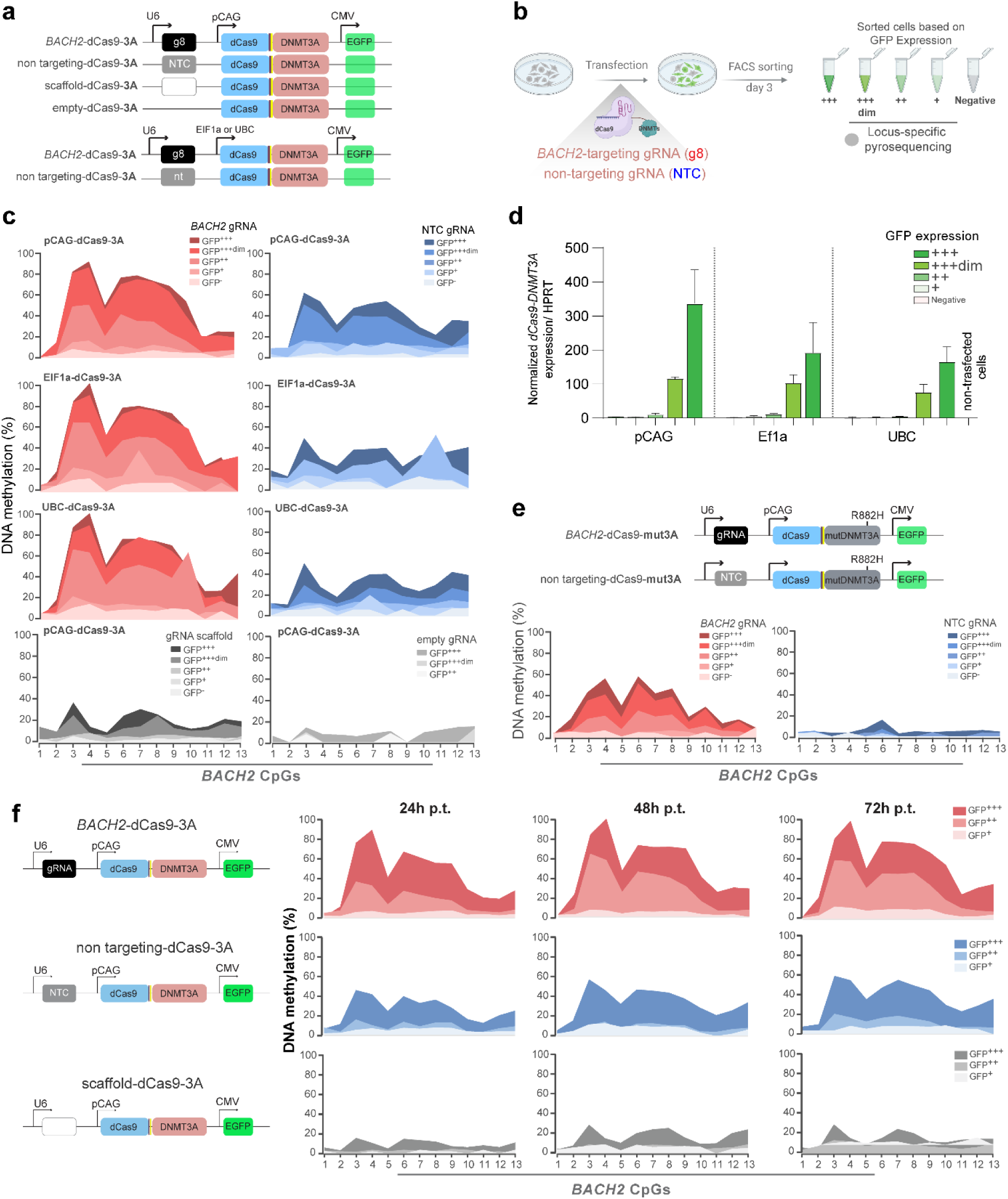
Factors influencing on-target methylation editing. **a.** Schematic of the designed. **b.** Experimental design where HEK293T cells were transfected with dCas9-DNMT3A-EGFP plasmids encoding *BACH2*-targeting gRNA8 (*BACH2*-dCas9-3A), non-targeting control gRNA (non-targeting-dCas9-3A), gRNA scaffold (scaffold-dCas9-3A) or no gRNA scaffold (empty-dCas9-3A). Successfully transfected cells were sorted 3 days post-transfection (p.t.) based on different GFP-intensity using fluorescent activated cell sorting (FACS). **c.** Impact of expression levels of dCas9-DNMT3A, based on sorting cells with varying levels of GFP intensity, in combination with strong (pCAG), moderate (EF1a), or weak (UBC) promoters, on methylation deposition induced by *BACH2* gRNA, NTC gRNA, gRNA scaffold or no gRNA scaffold. **d.** The expression levels of *dCas9-DNMT3A* under promoters with varying strengths assessed across samples with different GFP mean fluorescent intensities. **e.** Methylation deposition by dCas9-epimodifier comprising the catalytic domain of DNMT3A harboring the R882H mutation (dCas9-mut3A). **f.** Methylation deposition following transfection with *BACH2*-targeting gRNA8 (*BACH2*-dCas9-3A), non-targeting control gRNA (non-targeting-dCas9-3A) and gRNA scaffold (scaffold-dCas9-3A) 24, 48 and 72 hours (h) p.t..

Thus, to further investigate the impact of the expression levels of epimodifiers, we used dCas9-3A construct under a strong (pCAG), moderate (EF1a), or weak (UBC) promoter (**Fig. 2a**). The findings indicate that reducing the expression level of dCas9-3A, as reflected by different GFP mean fluorescent intensity and *dCas9-DNMT3A* transcripts levels (**Fig. 2d**), leads to a decrease in both on-target specific (g8) and unspecific (NTC) effects on *BACH2* locus, suggesting a correlation between the expression level and unwanted methylation deposition (**Fig. 2a-c**).

Our data revealed that transient transfection of the epimodifiers resulted in widespread gRNA sequence-independent methylation, indicating robust global activity of epimodifiers. To control the DNMT3A activity, we generated an additional DNMT3A variant harboring a R882H mutation (dCas-mut3A, **Fig. 2e**) in the catalytic domain affecting the enzymatic activity, CpG specificity and sequence preference^23^. The R882H mutation is a hotspot in acute myeloid leukemia (AML), leading to aberrant DNA methylation^24^. As demonstrated previously, this mutation resulted in a 50% reduction in the methylation efficiency of dCas9-mut3A, accompanied by a significant decrease in off-target effects, further emphasizing the important role of DNMT3A activity on the level in off-target effects^23^ (**Fig. 2e**). Of note, the data presented so far were collected from cells harvested 3 days p.t. and we therefore addressed the potential impact of shorter timeframes by harvesting cells after 24, 48, and 72 hours. Our data demonstrated that the on-target g8-guided methylation deposition at the *BACH2* locus increased rapidly to 90% within 24 hours, reached 100% by 48 hours, and remained stable at 72 hours (**Fig. 2f**). In parallel, the unwanted NTC-driven off-target methylation rose from 45% at 24 hours to 60% at 72 hours. A similar trend of methylation deposition over time was observed using the scaffold cassette. These findings suggest a pattern of increasing methylation deposition over time, indicating that methylation deposition is initiated rapidly following transfection.

Thus, the activity of dCas9-based epimodifiers is conditioned by the formation of a gRNA-dCas9 complex, with methylation deposition initiating rapidly following transfection. The level of unspecific effects, as seen with the NTC-driven methylation, is dependent on the amount of available epimodifier.

### Shared and distinct early impact of epimodifiers on transcriptome

We next investigated the functional impact of epimodifiers on gene expression by conducting RNA-seq analysis 3 days p.t. (**Suppl. Table 5**). Hierarchical clustering and Principal Component Analysis (PCA) indicated a clear separation according to the gRNA, with NTC interestingly exerting a greater variation compared to g8 (**Fig. 3a**). Overall, all epimodifiers with either g8 or NTC induced widespread but minimal transcriptomic alterations, i.e. less than 10% of the differentially expressed genes (DEGs defined as |log_2_FC| > 1) displayed |log_2_FC| > 2, compared to dCas9-d3A control (**Fig. 3b, c**). The strongest changes were observed with NTC, particularly in combination with dCas9-mut3A and CRISPRoff, resulting in 736 and 524 DEGs, respectively, compared to dCas9-d3A (**Fig. 3c**). The transcriptomic impact was milder upon g8 delivery in general, with the corresponding number of 181 and 153 DEGs and the highest number of 271 DEGs following dCas9-3A3L (**Fig. 3c**). The directionality of the changes did not appear skewed in most conditions with the exception of NTC-expressing dCas9-mut3A with 88% of the DEGs being upregulated (**Fig. 3c**). The majority of the DEGs displayed gRNA-specific pattern, i.e., with little overlap between g8 and NTC, and a large fraction of the differences (on average 72%) induced by one construct were shared with at least another epimodifier for a given gRNA (**Fig. 3c**, **Suppl. Fig. 3b**). Thus, all epimodifiers moderately affect the transcriptome in a gRNA-specific manner, with NTC-guided CRISPRoff and dCas9-mut3A exerting the largest influence three days after transient transfection.

**Figure 3.**
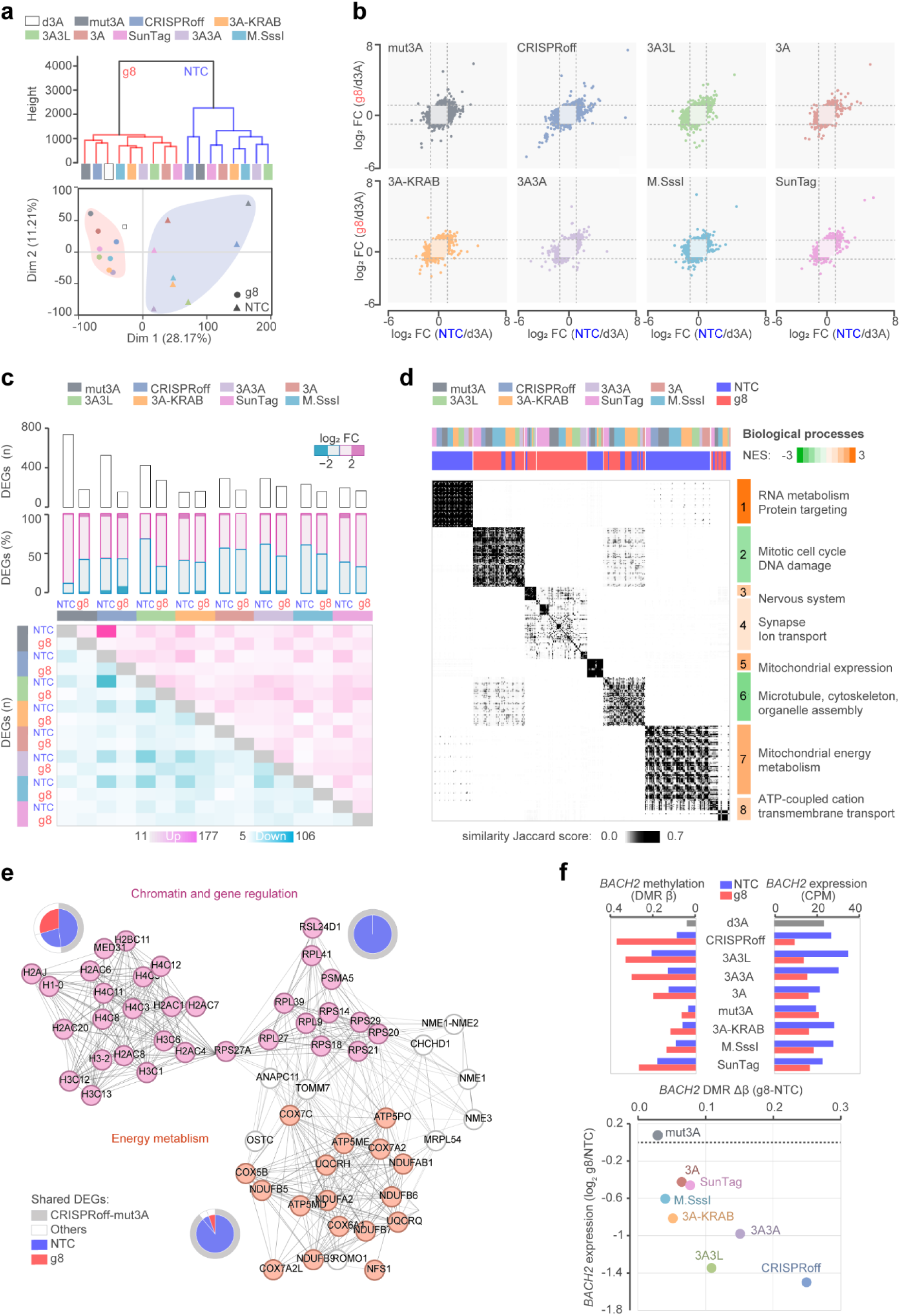
Early transcriptomic changes induced by the dCas9-epimodifiers. **a**. Hierarchical clustering and principal component analysis of RNA-sequencing data 3 days post-transfection with the dCas9-epimodifiers (3A, 3A3L, 3A3A, 3A-KRAB, mut3A, M.SssI, SunTag) or CRISPRoff (co-transfected with a gRNA-encoding plasmid). **b**. Correlation between transcriptional changes driven by *BACH2*-targeting (g8) and non-targeting control (NTC) gRNAs in comparison to the control deactivated DNMT3A (d3A) with the blue and pink colors reflecting downregulated and upregulated transcripts, respectively, between g8 and NTC. **c**. Total number of differentially expressed genes (DEGs, upper graph) and proportion in percentage (middle graph) of upregulated (pink color) and downregulated (blue color) DEGs between each dCas9-epimodifier and control d3A with |log_2_FC| > 1. The number of DEGs shared with at least one other epimodifier is detailed in the lower pairwise sharedness plot (the darker the color the highest number of genes being shared). **d**. Clustering of gene ontology terms, using GeneSetCluster tool, derived from the transcriptome of each epimodifier expressing either g8 or NTC compared to the control d3A, the Jaccard score depicted in grey gradient represents the degree of gene sharedness between sets (the darker the colors, the higher the number of genes being shared). The normalized enrichment scores (NES) illustrating upregulation and downregulation are indicated in orange and green colors, respectively for each cluster. **e**. STRING network of the core interconnected genes shared between epimodifiers identified using Density-Based Spatial Clustering of Applications with Noise algorithm. Pie charts summarize the dCas9-epimodifiers involved in the gene overlap for each sub-network, distinguishing the fraction shared between CRISPRoff and dCas9-mut3A from other shared genes. **f**. *BACH2* promoter differential methylation region (DMR, mean of CpGs β values) and gene expression (in counts per million, CPM, upper graph) and correlation between differences at *BACH2* promoter methylation (Δβ_g8-NTC_) and gene expression (log_2_ g8/NTC) between g8 and NTC gRNAs (lower graph).

To explore the potential biological relevance of the observed transcriptomic changes, we conducted Gene Set Enrichment Analysis (GSEA). Unsurprisingly, enrichment of terms in *Biological Processes* mirrored the impact of a given gRNA rather than the epimodifier (**Fig. 3d, Suppl. Table 6**). Indeed, NTC delivery resulted in a consistent upregulation (NES > 1.9, FDR < 0.05) of genes related to ‘*RNA metabolism’* and ‘*Protein targeting to endoplasmic reticulum’* (cluster 1) as well as ‘*Mitochondrial gene expression and energy metabolism’* (cluster 5 and 7) (**Fig. 3d**). NTC and to a lesser extent g8 induced genes involved in *ATP-dependent cation transport* (cluster 8). In contrast to the noticeable effect of NTC, g8-specific changes seemed modest and linked to the nervous system and ion transport (clusters 3 and 4, NES = 1.6 - 2.1, only GO:0031644 *‘Regulation of nervous system process’* for dCas9-3A3L passed FDR < 0.05) (**Fig. 3d**, **Suppl. Table 6**). The investigation of *Pathways* confirmed the drastic activation of gene regulation and energy metabolism, the core molecules forming an interconnected network of genes encoding multiple subunits of histones, ribosomes and mitochondrial enzymes, particularly following delivery of NTC with dCas9-mut3A and CRISPRoff (**Fig. 3e, Suppl. Table 6**).

At the on-target locus, the methylation editing efficiency at the *BACH2* promoter (**Fig. 1**) was reflected by varying degree of *BACH2* gene repression (**Fig. 3f**). The most potent epimodifiers, CRISPRoff, dCas9-3A3L and dCas9-3A3A, also exhibited the strongest *BACH2* repressing capacity (**Fig. 3f**) and *BACH2* gene was among the most downregulated transcripts by g8 in comparison to NTC for CRISPRoff (ranked 71 out of 263 downregulated transcripts) and dCas9-3A3L (ranked 28 out of 71 downregulated transcripts) (**Suppl. Table 5**).

Altogether, these data indicate that shortly after delivery all epimodifiers induce minimal but widespread transcriptomic changes that are shared by several constructs in a gRNA-specific manner. Remarkably, the non-targeting control gRNA exhibited a larger impact than g8. Such effect is particularly sizeable with CRISPRoff and dCas9-mut3A and reflects an upregulation of genes encoding histone subunits, ribosomal proteins and mitochondrial enzymes.

### Stability of on-target DNA methylation editing over time

To assess the stability of the deposited methylation at the *BACH2* promoter, we sorted GFP-positive transfected cells on day 3 p.t., replated them and harvested on days 3, 7, 14, 21 and 30 days p.t. for analysis using TMS **(Fig. 4a, Suppl. Table 7)**. The outcome of TMS, validated by pyrosequencing (**Suppl. Fig. 4**), confirmed the potent editing ability of multimerized epimodifiers 3 days p.t. using g8, with varying levels of unintended induction of differentially methylated regions (DMRs) at *BACH2* following NTC delivery (**Fig. 4b**). A residual effect of NTC remained until day 7 p.t. (average DMR β_NTC_ < 10%) and then decreased to control levels for all epimodifiers. On the contrary, the specific g8-induced methylation with CRISPRoff and dCas9-3A3L persisted (average DMR β_g8_ > 10%) until day 30 p.t. Furthermore, we found that the CpG sites targeted by the dCas9-gRNA complex remain strongly protected from DNA methylation until day 30 p.t., despite the transient nature of the transfection (**Fig. 4b**). The editing by other epimodifiers was less stable as methylation remained until 7 (dCas9-3A and SunTag) or 21 (dCas9-3A3A) days p.t. Accordingly, the analysis of the net specific effect of g8 (Δβ_g8-NTC_) at the *BACH2* promoter revealed CRISPRoff, dCas9-3A3L and dCas9-3A3A as the constructs most efficient in inducing stable methyl-deposition.

**Figure 4.**
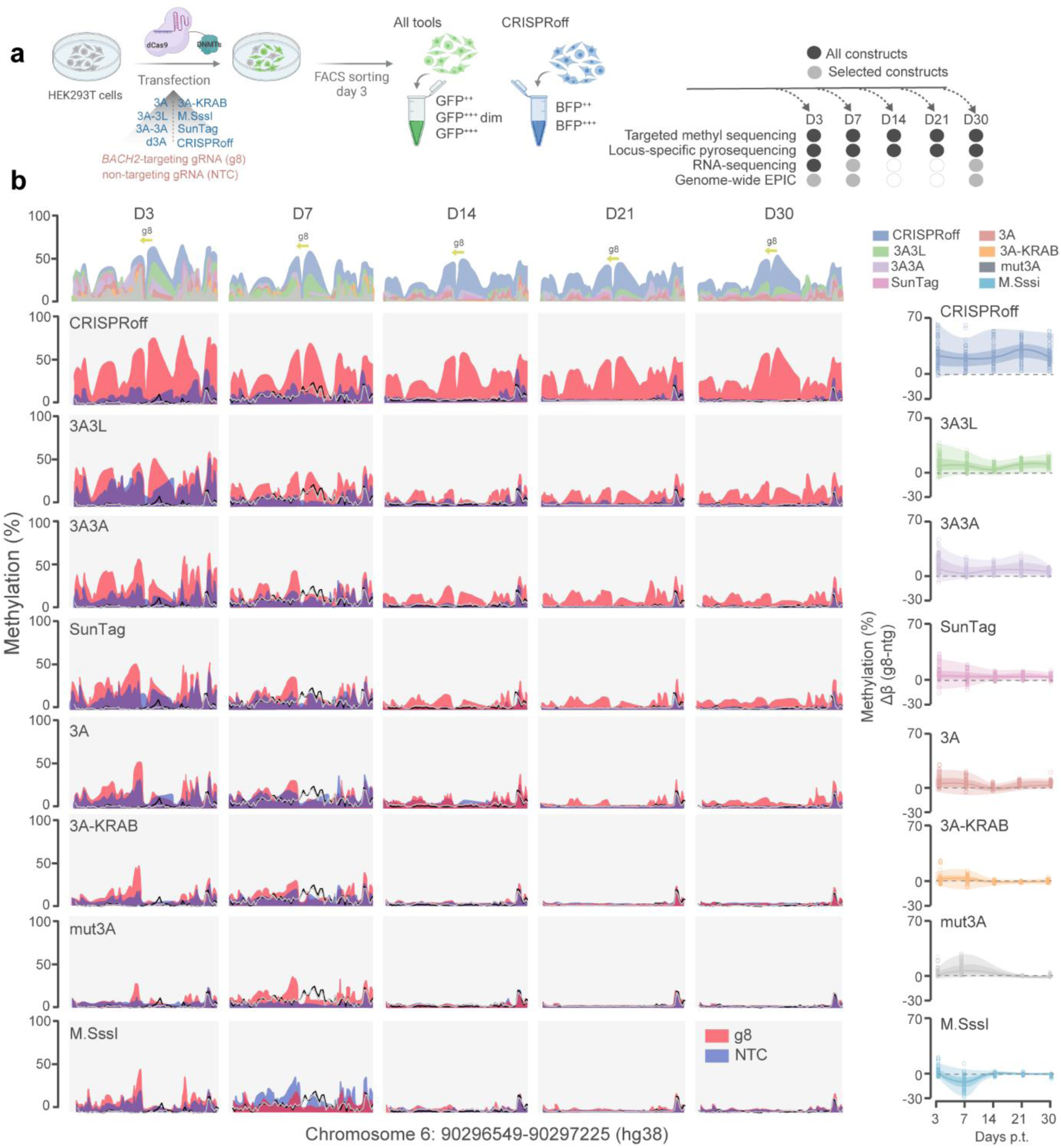
Stability of the on-target methylation editing over time. **a.** Experimental design where HEK293T cells were transfected with plasmids encoding gRNA-dCas9-epimodifier-EGFP (3A, 3A3L, 3A3A, 3A-KRAB, mut3A, M.SssI, SunTag) or CRISPRoff (co-transfected with a gRNA-encoding plasmid). Successfully transfected cells were sorted based on GFP or BFP signal 3 days post-transfection (p.t.) using fluorescent activated cell sorting (FACS), GFP/BFP positive cells were cultured and harvested at different time points for methylation and expression analyses. **b.** Methylation of CpGs at *BACH2* promoter 3 (D3), 7 (D7), 14 (D14), 21 (D21) and 30 (D30) days p.t. with the epimodifiers expressing either *BACH2*-targeting gRNA8 (g8) or non-targeting gRNA control (NTC). Yellow arrow indicated position of *BACH2*-targeting gRNA (g8). Methylation levels in the control conditions including cells expressing dCas9-deactivated DNMT3A (d3A) or non-transfected cells (NT) are depicted by the black and grey lines, respectively. Comparison of g8-driven methylation levels for all constructs per time point is illustrated by different turquoise blue colors. The net effect of g8 in comparison to NTC is represented on the right panel with the methylation differences of the differentially methylated CpGs (circles) centered on median (line), 25th and 75th percentile bounds (darker color) and minimum and maximum extending to the lowest / highest values (light color). Methylation data, assessed by targeted methylation sequencing, are available in Supplementary Tables 3 & 4.

### Genome-wide impact of epimodifiers on methylome over time

We further examined potential unspecific effects by profiling DNA methylation genome-wide using the Illumina Infinium EPIC array, focusing on the most efficient epimodifiers, i.e., CRISPRoff, dCas9-3A3L and dCas9-3A3A at day 3, 7 and 30 p.t. (**Fig. 5**). The conventional dCas9-3A and the mutated dCas9-mut3A were also included, along with the control deactivated dCas9-d3A. To mitigate the probability of identifying spurious alterations when analyzing > 850 000 CpGs, we prioritized DMR analysis with regions presenting coherent (same direction) and sizeable (|Δβ| > 0.1) methylation changes at several consecutive CpGs (see Methods). We identified a total of 34,785 DMRs across all epimodifiers and time points (**Table 1**, **Suppl. Table 8**).

**Figure 5.**
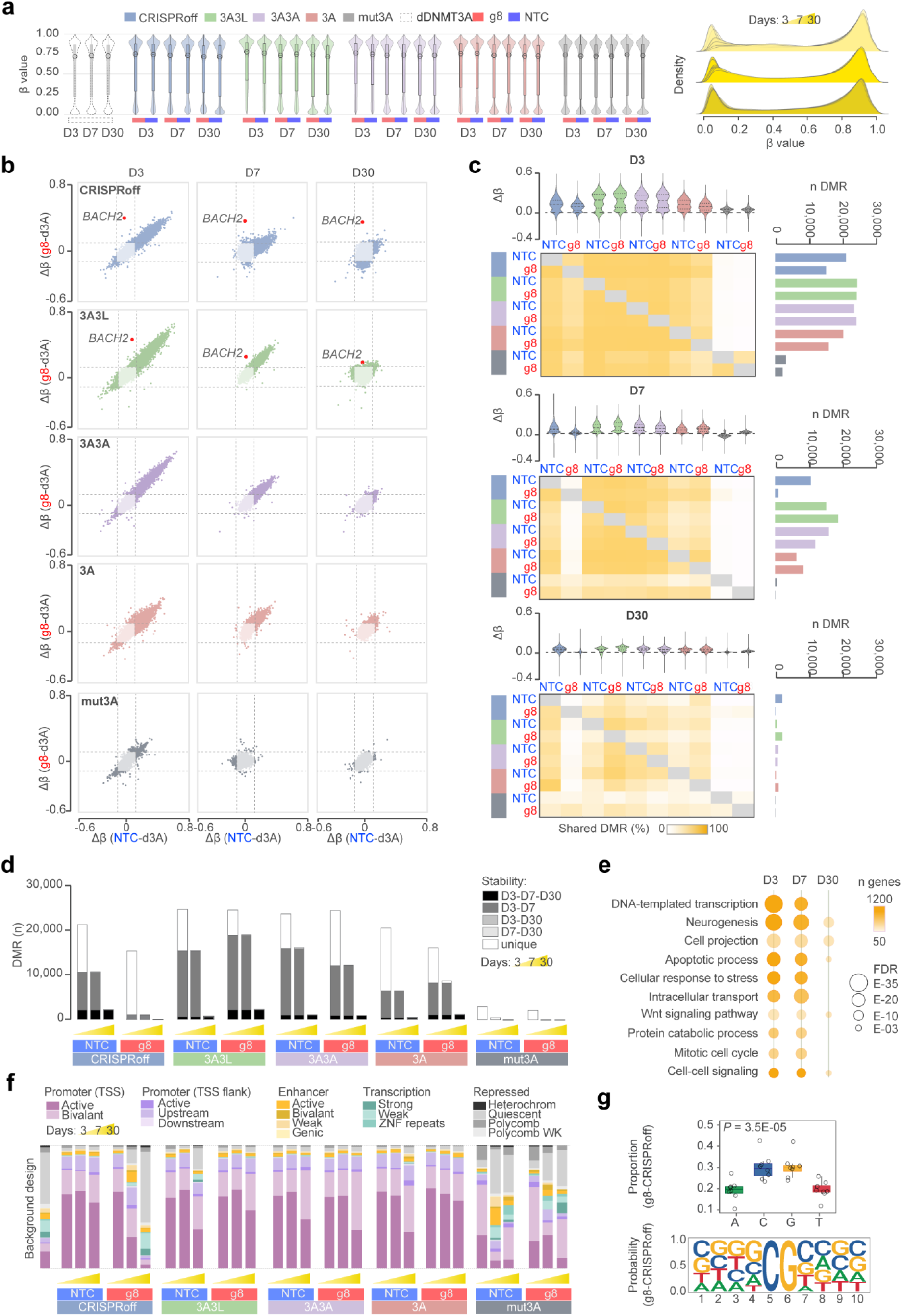
Genome-wide methylation induced by the four most potent dCas9-epimodifiers over time. **a.** Violin plot showing methylation level distributions across sample groups for day 3 (D3), 7 (D7) and 30 (D30) post transfection (p.t.). Overlayed boxplots indicate the interquartile range, and open circles mark the median values. Density plot is visualizing the overall distribution of methylation levels across all samples. **b.** Correlation between methylation changes at differentially methylated regions (DMRs) driven by *BACH2*-targeting gRNA8 (g8) or non-targeting control gRNA (NTC) in comparison to the control deactivated DNMT3A (d3A) at each time point, with the blue and purple colors reflecting hypomethylated and hypermethylated loci, respectively, between g8 and NTC. **c**. Effect size (violin plot) and total number (horizontal bar plot) of the DMRs induced by each dCas9-epimodifier at each time point. The fraction (%) of the DMRs that overlap between two conditions is depicted by the pairwise sharedness plot (the darker the yellow color the higher percentage of genes being shared). **d**. Fraction of DMRs stably altered between time points (gradient of grey colors) or specifically found only at day 3, 7 or 30 p.t. (white color). **e.** Top ‘Biological Processes’ gene ontology terms obtained using overrepresentation analysis of the DMR-genes that are shared between dCas9-epimodifiers. The circle size and color reflect the FDR significance and number of genes, respectively. **f.** Distribution of the DMRs according to chromatin state segmentation by hidden Markov model (ChromHMM) in HEK293T cells obtained from International Human Epigenome Consortium. The methylation data, assessed by EPIC array, is available in Supplementary Table 8. The statistics accompanying ChromHMM enrichment are available in Supplementary Table 9. TSS, transcription start site, Heterochrom, heterochromatin. **g.** Weblogo representation of sequence preferences for CRISPRoff gRNA8, showing GC enrichment, particularly in GCGC motifs. ANOVA – Tukey–Kramer’s test (sig P-value p < 0.05).

All epimodifiers induced extensive off-target methylation deposition compared to the control dCas9-d3A, particularly early after transfection and at low-methylated regions (β < 0.1) (**Fig. 5a,b**, **Table 1**), corroborated by observations at the single CpG level (**Suppl. Fig. 5 and 6**). Interestingly, the vast majority of the early changes (on average 89 ± 11% of the DMRs at day 3 p.t.) were shared between g8 and NTC and across all constructs (**Fig. 5b**). The dCas9-mut3A delivery induced fewer and smaller changes that differed from the other by their binomial distribution at both low and high methylation levels (**Fig. 5a,b**, **Table 1**, **Suppl. Fig. 5**). The number and amplitude of genome-wide changes markedly decreased with time in all conditions, with off-target changes induced by g8-guided CRISPRoff drastically subsiding already at day 7 p.t. (**Fig. 5b**, **Table 1**).

**Figure 6.**
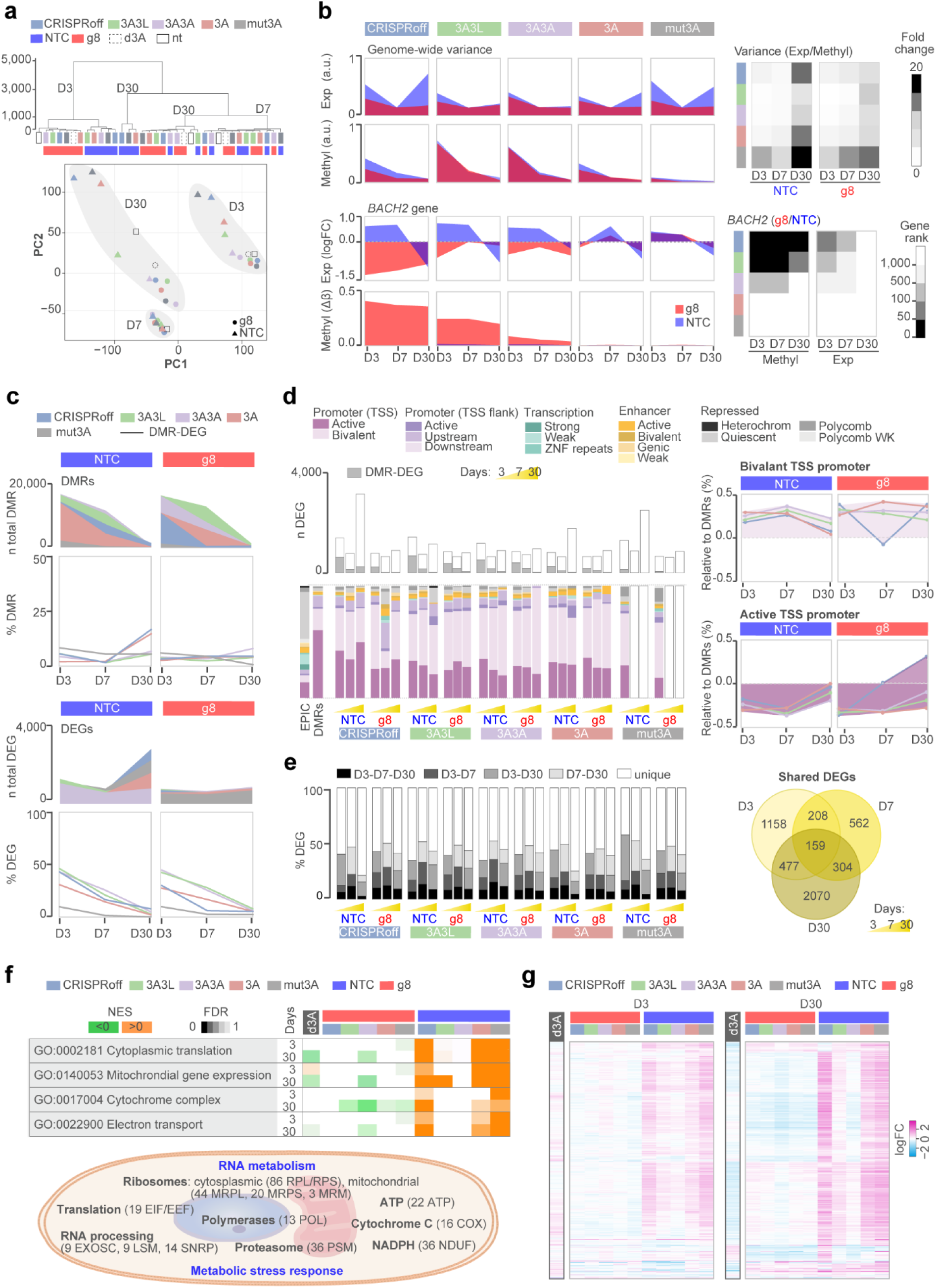
Transcriptional changes induced by the most potent dCas9-epimodifiers over time. **a**. Hierarchical clustering and principal component analysis of RNA-sequencing data. **b**. Genome-wide and on-target (*BACH2*) methylation and transcriptome effects. The top and bottom heatmaps represent the fold change of expression/methylation variances (the greater, the darker the gradient) and the rank of *BACH2* differentially methylated region (DMR) and gene among all DMRs and the downregulated genes (the darker the more highly ranked), respectively. **c**. Number of total DMRs (|Δβ| > 0.1) and differentially expressed genes (DEGs) (|log_2_ FC| > 1) between a given gRNA-epimodifier and the control dCas9-d3A condition and the percentage of them displaying both methylation and expression changes (DMR-DEGs). **d**. Number of DEGs (|log_2_ FC| > 1) and DMR-DEGs (grey color) and distribution of the DMR-DEGs according to chromatin state segmentation by hidden Markov model (ChromHMM) from HEK293T cells (obtained from International Human Epigenome Consortium). The EPIC background and average DMRs at day 3 post-transfection (p.t.) are shown as references. Detailed enrichment for bivalent and active TSS promoter features (in percentage increase or decrease) of DMR-DEGs compared to all DMRs for each epimodifier. The statistics accompanying ChromHMM enrichment are available in Supplementary Table 9. TSS, transcription start site, heterochrom, heterochromatin. **e**. Proportions of DEGs that overlap (or not) between time points. The Venn diagram represents the number of stable or transient DEGs that are shared by at least two epimodifiers. **f**. Top Gene Set Enrichment Analysis (GSEA) terms with green/orange colors representing negative/positive normalized enrichment scores (NES) and color gradient reflecting the FDR significance. The schematic illustrates the main processes and families of genes implicated by the most significant GSEA terms. **g**. Heatmap of the 647 DEGs corresponding to the top GSEA terms with upregulated and downregulated genes depicted in pink and blue, respectively. Results arising from the effect of transfection with the control dCas9-dDNMT3A (d3A) compared to non-transfected cells are highlighted in black.

Specific on-target methylation changes at the *BACH2* promoter confirmed the outcomes of TMS (**Fig. 4b**). *BACH2* methylation deposition was apparent at all time points after delivery of both CRISPRoff (Δβ_g8-d3A_ = 0.35-0.40) and dCas9-3A3L (Δβ_g8-d3A_ = 0.19-0.24) (highlighted in **Fig. 5b**). For dCas9-3A3A the average methylation deposition was below the threshold for dCas9-3A3A (Δβ_g8-d3A_ = 0.03-0.08) and could not be detected in dCas9-3A and dCas9-mut3A at any time. Of note, only 8 out of 55 CpGs (detected by TMS) in the *BACH2* DMR are included in the array design causing underestimation of the on-target effects with EPIC compared to TMS.

We next asked to what extent the off-target DMRs detected 7 and 30 days following transfection resulted from early changes that remained stable or from secondary effects. A large fraction of the changes (average [range] = 55% [30-76] across epimodifiers) induced at day 3 p.t. remained stable for at least one week for all constructs except dCas9-mut3A and g8-guided CRISPRoff (**Fig. 5c**, **Suppl. Fig. 7**) and nearly 10% (2,000 DMRs) of the early changes induced by NTC-expressing CRISPRoff persisted at day 30 p.t. (**Fig. 5c**, **Suppl. Fig. 7**). Consistent with the low number of DMRs 3 days p.t., specific g8-guided CRISPRoff displayed a distinctive non-overlapping methylation pattern with a particularly low fraction of early off-target effects persisting for 7 (5%, 832/16107) and 30 days (0.3%, 51/16107) (**Fig. 5b,c**). Functional annotation of the genes that associated with the DMRs shared between epimodifiers reflected the substantial overlap between 3 and 7 days p.t. (**Fig. 5d**) and indicated a strong enrichment of GO terms pertaining to transcription (e.g., 1,158 genes involved in DNA-templated transcription associate with DMRs at day 3 p.t.) followed by, e.g., response to stress, apoptotic/mitotic processes and cell-cell signaling (**Fig. 5e**). Similar findings were obtained after GO analysis of the DMRs that persisted for one month (**Suppl. Table 9**).

We addressed the genomic properties of the off-target DMRs according to chromatin regulatory features in HEK293T cells annotated by International Human Epigenome Consortium (IHEC). The DMRs induced by CRISPRoff, dCas9-3A3L, dCas9-3A3A and dCas9-3A at day 3 p.t. were significantly enriched (*P*-value < E-15) in active promoter-like features and depleted in other regulatory features such as enhancers, transcribed and repressed regions (**Fig. 5f, Suppl. Table 10**, **Suppl. Fig. 7**). This preferential enrichment in active promoters prevailed until day 30 p.t. for dCas9-3A3L, dCas9-3A3A and dCas9-3A (*P*-value < E-15) irrespective of the gRNA (**Fig. 5f, Suppl. Table 10**, **Suppl. Fig. 7**). In contrast, the enrichment of changes resulting from g8-guided CRISPRoff editing shifted from active promoters at day 3 p.t. to active enhancers 7 days p.t. (*P*-value < E-15) and actively transcribed regions 30 days p.t. (*P*-value = 2E-09). Notably, dCas9-mut3A displayed off-target effects distinct from other epimodifiers with significant enrichment of DMRs in both bivalent promoters and enhancers as well as Polycomb-repressed regions at day 3 p.t. (*P*-value < E-15) followed by a shift to enhancer-like features at day 7 p.t. (NTC, *P*-value < E-15) and zinc-finger transcription factor (ZNF) gene repeats at day 30 p.t. (*P*-value = 2E-07) (**Fig. 5f**, **Suppl. Table 10**, **Suppl. Fig. 8**). Similar results were observed when single CpGs were analyzed (**Suppl. Table 10**). Consistent with the enrichment of DMRs at promoters across conditions, sequence preference analysis indicated that methylation occurs at GC-rich sequences, particularly in GCGC motifs (exemplified with CRISPRoff-g8 at day 3 in **Fig. 5g**), with no difference observed between gRNAs, timepoints or epimodifiers (**Suppl. Table 11**).

Thus, the most potent epimodifiers display noticeable gRNA independent and largely overlapping genome-wide off-target effects that are enriched in active GC-rich promoter-like features and a fraction of which remains stable over time. Notably, the specific gRNA-guided CRISPRoff induces potent and stable on-target methylation with fewer, milder and less stable off-target alterations.

### Genome-wide impact of epimodifiers on transcriptome over time

To delineate the potential functional impact of the off-target methylation, we investigated transcriptional changes in the same samples with RNA-sequencing. Overall, the samples clustered primarily according to the time point followed by the gRNA (**Fig. 6a, Suppl. Table 12**). The genome-wide variance indicated that NTC produced greater heterogeneity at all time points compared to g8 gRNA (**Fig. 6b, Suppl. Fig. 9**). CRISPRoff, dCas9-mut3A and dCas9-3A elicited high NTC-driven transcriptomic heterogeneity and moderate methylome variance at days 3 and 30 p.t., e.g., leading to 10- to 20-fold higher transcriptome vs. methylome variability at day 30 (**Fig. 6b, Suppl. Table 13**). On the contrary, dCas9-3A3L and dCas9-3A3A produced a strong early unspecific methylome effect and mild transcriptomic variability both with NTC or g8, this heterogeneity subsided over time (**Fig. 6b**).

We addressed the methylation-expression correspondence using CpG-gene correlation analysis (Spearman *P* < 0.05, **Suppl. Table 14, Suppl. Fig. 10**) and examined the genes displaying both methylation (DMRs with |Δβ| > 0.1) and expression (DEGs with |logFC| > 1) changes, referred to as DMR-DEGs. Only a minor fraction (< 17%) of the gene-mapping DMRs resulted in transcriptional differences at any given time (**Fig. 6c**). On the other hand, on average 40% of DEGs exhibited methylation changes 3 days p.t. and this proportion decreased to 20% after 7 days and further to 4% after 30 days p.t. (**Fig. 6c**, **Suppl. Table 14**). The DMR-DEGs were generally remarkably enriched in promoters and depleted in repressed chromatin and mapped preferentially to bivalent (as opposed to active) TSS promoters and to bivalent and active enhancers, compared to all DMRs (**Fig. 6d**, **Suppl. Table 10**). The DMR-DEGs displayed the expected predominant promoter hypermethylation but not the canonical relationship between promoter and enhancer methylation and gene repression since genes displayed both increased and decreased expression (**Suppl. Table 10**), as seen for all CpG-gene pairs (**Suppl. Fig. 10**). In stark contrast to all DEGs, the majority of the DMR-DEGs (on average 74% across epimodifiers) appeared unique to a given time point, the remaining (1-14%) overlapped between day 3 and 7 p.t. (**Suppl. Table 14**), mirroring the pattern observed with all DMRs (**Fig. 5c**).

Nearly half of the DEGs overlapped between time points, particularly between days 3 and 30 p.t. on average 9% of the DEGs persisted throughout 30 days (**Fig. 6d**, **Suppl. Fig. 11)**. The overlap between early (day 3) and late (day 30) changes was notably higher for NTC guided dCas9-mut3A (41%), CRISPRoff (28%), and dCas9-3A (24%), compared to other gRNA-epimodifier pairs (average 18%) (**Fig. 6e**, **Suppl. Fig. 11**). We addressed the biological relevance of the seemingly independent impact on methylome and transcriptome using GSEA (**Suppl. Fig. 11, Suppl. Table 15**). Consistent with the gene overlap (**Suppl. Fig. 10**), most of the perturbations were shared by constructs expressing either g8 or NTC and overlapped between day 3 and 30 p.t., underscoring temporally concordant gRNA-specific effects (**Suppl. Fig. 11**). The changes induced by g8-expressing epimodifiers resulted in very few significant GO terms related to upregulated nervous and immune system processes (**Suppl. Table 15**). Interestingly, terms pertaining to nervous system processes were the only GO terms found significantly enriched for DMR-DEGs, irrespective of the gRNA, representing changes at genes encoding subunits of potassium, calcium channels and glutamate receptors involved in regulation of membrane potential and synaptic transmission (**Suppl. Table 15**). The biological cluster that reflected the largest disparity, both early and late, between gRNAs related to NTC-guided dCas9-mut3A, CRISPRoff, and dCas9-3A converged to activated RNA and energy metabolism (**Suppl. Fig. 11, Suppl. Table 15**). This is exemplified by enrichment (NES > 2, FDR < E-15) of terms referring to *‘Cytoplasmic translation’*, *‘Mitochondrial gene expression’*, ‘*Cytochrome complex assembly’* or *‘Electron transport chain’* (**Fig. 6f, Suppl. Table** 1**5**). The sustained and stronger transcriptional impact evoked by NTC delivery on translation and mitochondrial respiration could be confirmed when comparing g8 vs. NTC at the group level (**Suppl. Fig. 12**). Overall, the most significant GSEA terms resulted from joint, albeit mild (average logFC = 1 ± 0.9), dysregulation of 647 genes encoding multiple subunits of cytoplasmic and mitochondrial ribosomes, enzymes involved in translation and RNA processing (**Fig. 6g**, illustrated in **Fig. 6f, Suppl. Table 15**). Mobilization of transcripts implicated in gene expression was accompanied with upregulation of genes encoding subunits of complexes involved in energy demand, mitochondrial respiration, oxidative stress, and proteasomal degradation and, importantly, could not be accounted to the impact of transfection alone (**Fig. 6g, Suppl. Table 15**). Late dysregulation of the genes included in the top GSEA terms could not be explained by methylation differences at the corresponding genes for dCas9-3A and dCas9-mut3A while only modest methylation changes (average ± SD DMRs Δβ_NTC-d3A_ = 0.06 ± 0.03) could be detected at these DEGs for CRISPRoff (**Suppl. Fig. 11**), overall suggesting a methylation-independent origin.

To investigate the role of gRNA sequences in genome-wide off-target effects, we examined in-silico predicted off-target sites with up to 5 mismatches (see methods). The number of predicted off-targets varied across tools, with a notable proportion involving 5 mismatches, highlighting potential off-target effects even with high mismatch tolerance **(Suppl. Table 16)**. As expected, g8 had significantly more predicted off-targets than NTC (1637 vs. 333, respectively), which was designed to avoid genomic targeting. Importantly, only a small fraction of predicted off-targets displayed methylome (g8 = 0.002 %, NTC = 0.05 % showed |Δβ| > 0.05) and transcriptome (g8 = 0.05 %, NTC = 0.07 % showed |logFC| > 0.05) changes on day 3 p.t. **(Suppl. Table 16**), unlinking gRNA-binding with observed off-target effects.

Thus, the expression of non-targeting control gRNA induced pervasive, mostly methylation-independent, transcriptional changes in genes involved in RNA and energy metabolism, likely reflecting the cellular adaptations in response to genomic and oxidative stress. Such reprograming could be detected one month after transfection and was more pronounced in association with CRISPRoff and dCas9-mut3A.

### On-target correlation of methylation and expression

We further explored on-target effect and conducted ‘Upstream Regulators’ analysis, using Ingenuity Pathway Analysis, of the DEGs induced by the two most potent *BACH2*-repressing epimodifiers, i.e. CRISPRoff and dCas9-3A3L. Analysis of the DEGs induced by g8-expressing CRISPRoff three days p.t. identified *BACH2* as ‘Master regulator’ of a ‘Causal network’ that includes *BACH2*, *PRDM1*, *IRF4*, *PRF1*, *TBX21*, *STAT4* and *GZMB* co-regulated genes (**Fig. 7a**).This *BACH2*-centered module being further embedded into a larger immune-related causal network and among the closest interactors, *IRF4* and *PRDM1* displayed dynamic gene regulation over time, suggesting their involvement in shared regulatory networks and biological pathways (**Suppl. Table 15**).

**Figure 7.**
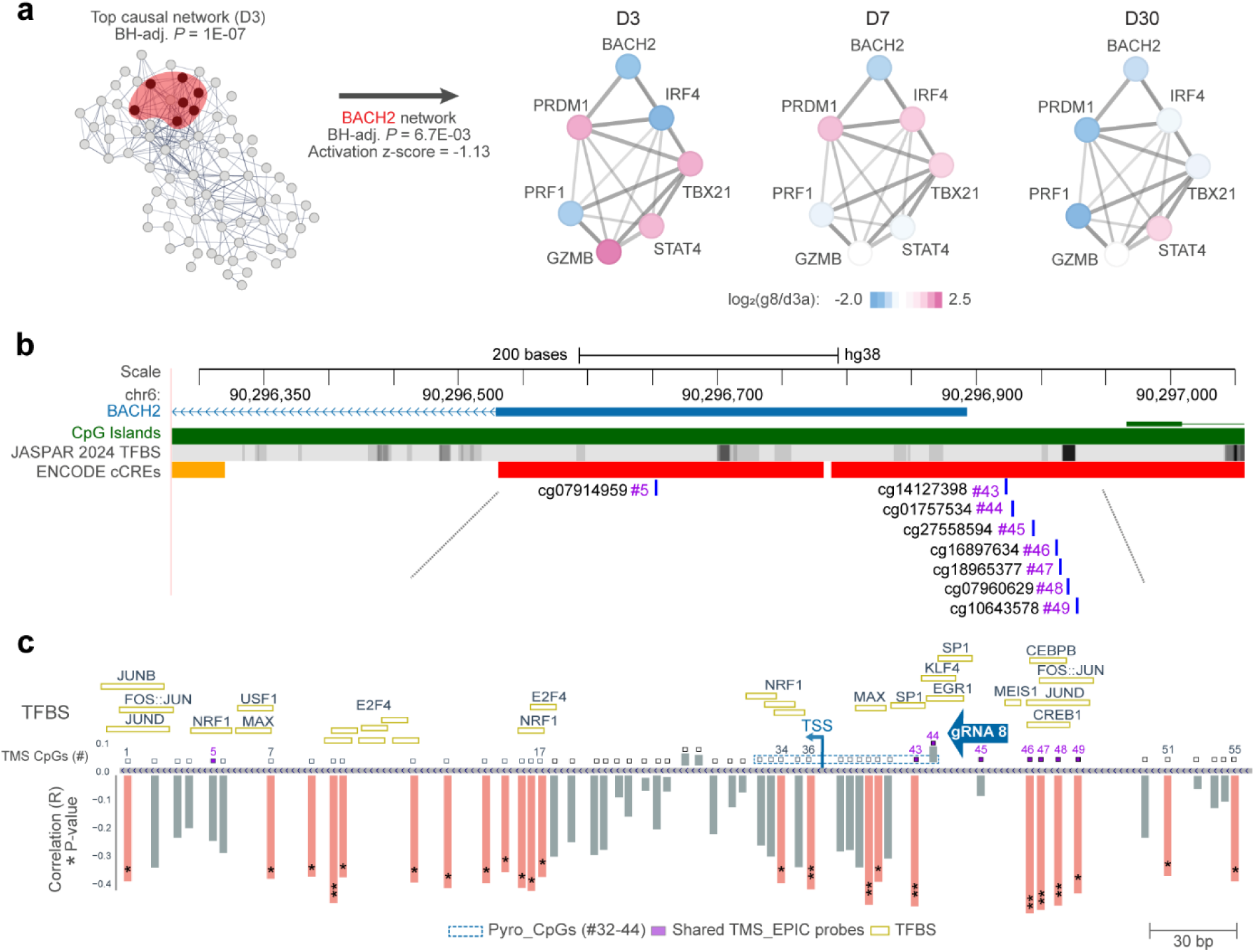
*BACH2* gene network and methylation control. **a.** Top significant causal network and detailed BACH2-master regulator network obtained using Ingenuity Pathway Analysis (Benjamini-Hochberg corrected P < 0.05) of RNA-sequencing data obtained 3 days post-transfection with CRISPRoff and *BACH2*-targeting gRNA 8 (g8), visualized using STRING with blue and pink colors reflecting downregulated and upregulated genes, respectively. **b.** A schematic of the *BACH2* locus illustrating the EPIC probes and regulatory features. **c.** Details of the CpG sites included in the targeted methylation sequencing design (CpGs 1-55) and pyrosequencing assay, along with the gRNA8 binding site and transcription factor binding sites (TFBS), with several methylation-sensitive TFs identified (orange rectangles). CpG identifiers in the TMS design are presented as sequential numbers (CpGs 1–55; see Supplementary Table 15), with Pearson correlation coefficient (R) values for correlation between methylation and *BACH2* expression indicated as bar plots on the map. Significant correlations are marked by light red bars, while non-significant ones are indicated in gray, with p-values represented as * and ** for p < 0.05 and p < 0.01, respectively.

To refine the identification of the CpG sites involved in methylation-mediated changes in *BACH2* gene expression **(Fig. 7b**, **Suppl. Table 17)**, we performed correlation analysis using TMS (including 55 CpGs at *BACH2*) and RNA-sequencing. The methylation levels of 23 CpGs located in two distinct blocks (CpGs 7-17 and CpGs 34-55) showed significant negative correlation (R = −0.34 to −0.49, Pearson p < 0.05) with *BACH2* expression **(Fig. 7c**, **Suppl. Table 17)**. Among the 8 CpGs shared between the TMS and EPIC designs (highlighted in purple in **Fig. 7c**), 5 probes located upstream and downstream of the gRNA8 binding site showed a significant negative correlation with *BACH2* expression. Interestingly, methylation patterns at CpG sites at and near the gRNA binding site (CpG 44-45) differed between the g8 and NTC gRNAs insofar NTC but not g8 induced undesired on-target methylation (**Suppl. Fig. 2**, **Suppl. Table 7)**. Transcription factor (TF) binding sequence analysis revealed multiple TF binding sites where MEIS1 and CREB1 had the highest binding scores **(Suppl. Table 18)**. Among all TFs, a notable subset (depicted in **Fig. 7b**) is methylation-sensitive^25,26^, underscoring their potential roles in the epigenetic regulation of *BACH2*. Of particular interest is the AP1 TF group, notably FOS, JUN, ATF, and MAF families, which are highly expressed in HEK293T cells^27^, bind to the sequence encompassing CpG sites, and show a high correlation with *BACH2* expression. The presence of these methylation-sensitive TFs highlights a complex interplay between DNA methylation and transcriptional regulation at the *BACH2* promoter, emphasizing their potential influence on gene expression dynamics.

## Discussion

Due to the diversity of CRISPR/dCas9-based strategies developed to manipulate DNA methylation, varying with respect to the utilized epimodifier, delivery method, target locus, cellular system and methylome coverage, it has been challenging to compare available systems. In this study, we expanded the existing toolbox with novel dCas9-based DNA methylation editing systems and systematically compared the existing epimodifiers under constant experimental conditions. Our work addresses a critical gap in the field by focusing on epimodifiers’ on-target efficacy and stability, and off-target activity at the methylome and transcriptome levels, as well as off-target mitigating strategies and factors influencing these outcomes.

Our data demonstrate that all tested epimodifiers effectively induce DNA methylation at the target *BACH2* promoter and that the deposition occurs rapidly, within 24 hours of transfection, consistent with previous findings using inducible dCas9-DNMT3A^18^. Multimerized variants, particularly CRISPRoff and dCas9-3A3L, exhibited the best methylating capability accompanied by robust transcriptional repression. The specificity of CRISPRoff-induced changes at *BACH2* was further reflected by the detection of *BACH2*-induced network of co-regulated genes. Interestingly, as previously shown for dCas9-3A3L^16^, we observed a long-lasting protection of CpG sites covered by the dCas9-epimodifiers despite the transient nature of transfection. This suggests a lasting footprint that persists even after the presence of epimodifier diminishes.

We further correlated *BACH2* expression following methylation deposition with the DNA methylation levels at single CpGs using robust targeted sequencing method for quantification. We found that among many differentially methylated CpG sites, only methylation at specific CpGs significantly correlates with gene expression. The overlap of these CpGs with the binding motifs of TFs, particularly those that are methylation-sensitive, further supports their gene regulatory role under the given conditions. Interestingly, while we observed increased methylation at the *BACH2* locus early after transfection with non-targeting gRNAs, there was no expected anti-correlation between methylation and gene expression, even when focusing on these specific regulatory CpGs. The underlying reasons remain to be ascertained and may suggest uncharacterized aspects of gene regulation. Thus, in addition to linking a genomic region with the regulation of a specific gene, methylation editing provides means to dissect the regulatory landscape of CpGs, advancing our understanding of methylation-mediated gene regulation. This is particularly relevant given the coordinated nature of methylation at adjacent CpGs^28,29^ that hampers characterization of the functionally relevant CpGs in regions that associate with various traits and diseases.

CRISPRoff emerged as the most effective epimodifier for depositing and maintaining on-target locus methylation during the tested 30 days post-transfection. The enhanced performance of CRISPRoff can be attributed to its design, particularly the inclusion of the KRAB domain, one of the most potent transcriptional repressors^20,30^. By interacting with its associated protein (KAP1), KRAB recruits chromatin-modifying complexes (e.g., HP1, HDACs, HTMs) leading to chromatin remodeling, histone modification, and the formation of facultative heterochromatin, contributing to stable DNA methylation and long-term transcriptional repression^31,32^. Importantly, this effect depends on the orientation of the KRAB domain and its positioning upstream of the dCas9-3A3L sequence, as previously demonstrated^21^ and further supported in our study by suboptimal performance of the 3A-KRAB configuration in terms of both efficacy and stability of DNA methylation. Notably, despite lacking the KRAB repressor domain, the dCas9-3A3L followed CRISPRoff in efficacy and displayed rapid comparable on-target methylation that persisted for 30 days. Thus, dCas9-3A3L, followed by newly designed dCas9-3A3A, provide alternatives when the effect of manipulating DNA methylation alone is the focus of investigation.

Besides the on-target effect at *BACH2*, we observed significant alterations in the methylome and transcriptome over time, reflecting both the transient and lasting impacts of the epimodifiers. Regions with low or intermediate baseline methylation, notably promoter-related GC-rich open chromatin, exhibited the highest methylation gains following delivery of both targeting- and non-targeting gRNAs. These observations align with studies showing that Cas9 activity is influenced by genomic context, including the host organism and cell type^18,37–39^. Importantly, while unintended methylation induced by non-targeting gRNA at the *BACH2* locus gradually diminished over time, specific gRNA-induced on-target methylation persisted. Inversely, genome-wide off-target methyl-deposition following specific gRNA rapidly subsided as opposed to off-target methylation induced by the specific non-targeting control gRNA. Off-target prediction analysis confirmed that unspecific effects cannot be directly linked to binding of gRNA-specific sequences at predicted loci, even when five mismatches were tolerated. One cannot exclude that biophysical parameters, including the sequence preference of dCas9 which differs from that of Cas9^27,28^ or the addition of editor proteins like DNMTs affecting the interaction between the dCas9 complex and the genome^40^, further complicate off-target prediction.

Nevertheless, the high methylation levels observed irrespective of the gRNA or even in the absence of a gRNA support substantial gRNA-independent activity which has also been noted in previous studies^17,18,41^. Notably, we found that the activity of epimodifiers is conditioned by the formation of a gRNA-dCas9 complex, with the amount of available epimodifier influencing unspecific methylation trends. Accordingly, the dCas9-epimodifier complex may engage in a global DNA search, preferentially binding to accessible chromatin and leading to off-target deposition. Thus, factors such as the process of Cas9 searching, overexpression of dCas9-epimodifier and chromatin accessibility significantly contribute to off-target effects^17,18,42–44^. The widespread off-target activity by the non-targeting control gRNA rendered the comparison with the targeting g8 gRNA uninformative but using deactivated DNMT3A as a control might provide a clearer baseline for assessing targeted methylation. Nevertheless, our results also reinforce the importance of dosage and time control in minimizing off-target effects and further encourage the implementation of strategies such as controlling the expression window and levels of epimodifiers as well as directing the complex to an irrelevant locus in the genome that would help mitigate off-target methylation.

We further tested the impact of DNMT3A catalytic activity on methylation editing by designing a new dCas9-mut3A epimodifier comprising the AML R882H mutation known to decrease enzymatic activity^23^. This led to a 50% reduction in the on-target methylation, accompanied by a significant decrease in off-target effects, further underscoring the important role of DNMT3A activity in regulating off-target effects. Moreover the fact that this mutation alters the enzymatic activity, CpG specificity, and sequence preference^23^, likely explains the distinct pattern of methylation changes observed by us and reported for other DNMT3A mutations by others^33,34^. Our data align with evidence indicating that R882H mutation stabilizes the formation of large oligomeric DNMT3A complexes, which are less enzymatically active and inhibit the function of wild-type DNMT3A^35,36^. Interestingly, despite its reduced methylation efficacy, dCas9-mut3A induced sizeable transcriptional changes, possibly indicating molecular exhaustion and potential cellular stress responses. These findings highlight the intricate interplay between methylation-dependent and independent regulatory mechanisms. Moreover, the observed changes may point to broader functional consequences of the R882H mutation extending beyond its role in DNA methylation.

Notably, the vast majority of the off-target methylation deposition, even though mapping to promoters, did not associate with expression changes. Inversely, the unspecific transcriptional noise observed early after delivery of non-targeting gRNA control and maintained over time, especially for CRISPRoff and dCas9-mut3A, is not mirrored by variation in methylome. The sustained methylation-independent impact on RNA processing and energy metabolism could reflect persistent cellular adaptations to overexpressing non-targeting short RNAs. More generally, the discrepancies between off-target promoter methyl-deposition and transcriptional effect suggest that additional mechanisms, such as baseline level of transcription, TF availability, chromatin modifications and subnuclear localization, might be at play in the transcriptional regulation. Interestingly, the differentially methylated regions that associated with alterations in gene expression preferentially mapped to bivalent chromatin states, defined as regulatory domains with both activating and repressive histone marks that promote poised gene state^45^. The dual directionality of the effect, with both activated and repressed gene transcription upon *de-novo* methylation of CpGs in bivalent regions is consistent with previous report^46^ and further highlights the regulatory potential of bivalent chromatin states in transcriptional regulation.

## Conclusions

This study provides a comprehensive evaluation of dCas9-based DNA methylation editors, identifying their strengths and limitations. While CRISPRoff demonstrates superior efficiency, stability and specificity, off-target activity remains a challenge. Our findings stress the importance of selecting appropriate controls and epimodifiers to minimize off-target effects in epigenetic studies. Further refinement of construct designs and gRNA strategies, alongside investigations of the molecular mechanisms underlying specific and stable methylation, will be essential for advancing the utility of DNA methylation editors in research and their therapeutic applications.

## Acknowledgements

We thank Dr. Alex Espinosa for his invaluable scientific advice and for providing the pEGFP-C1 plasmid. The computations were enabled by resources provided by the Swedish National Infrastructure for Computing (SNIC) through the Uppsala Multidisciplinary Center for Advanced Computational Science (UPPMAX), partially funded by the Swedish Research Council through grant agreement no. 2018-05973. The authors acknowledge support from the National Genomics Infrastructure in Stockholm funded by Science for Life Laboratory, the Knut and Alice Wallenberg Foundation, and the Swedish Research Council (through grant agreement no. 2018-05973). We thank QIAGEN for including us in the testing of the QIAseq Targeted Methyl Panels. We thank the Center for Molecular Medicine (CMM) for providing research facilities, including the KI Gene and Flow Cytometry Core, among others.

## Review history

The review history is available as Additional file X.

## Funding

This work was supported by grants from the Swedish Research Council, the Swedish Association for Persons with Neurological Disabilities, the Swedish Brain Foundation, the Swedish MS Foundation, the Stockholm County Council (ALF project), Karolinska Institutet and European Union Horizon 2020 Research and Innovation Programme/European Research Council Consolidator (grant Epi4MS, 818170). M. Pahlevan Kakhki was supported by the McDonald fellowship from the Multiple Sclerosis International Federation (MSIF) and ECTRIMS. This project has received funding from the Innovative Medicines Initiative 2 Joint Undertaking (JU) under grant agreement No 875510. The JU receives support from the European Union’s Horizon 2020 research and innovation programme and EFPIA and Ontario Institute for Cancer Research, Royal Institution for the Advancement of Learning McGill University, Kungliga Tekniska Hoegskolan, Diamond Light Source Limited. This communication reflects the views of the authors and the JU is not liable for any use that may be made of the information contained herein. G. Zheleznyakova was supported by the fellowship from the Swedish Society for Medical Research (SSMF). L. Kular was supported by fellowship from the Margaretha af Ugglas Foundation.

## Availability of data and materials

The sequencing data are available in Gene Expression Omnibus (GEO, XXX). All the plasmids are deposited in Addgene and are available under the Maja Jagodic’s Lab Materials (https://www.addgene.org/Maja_Jagodic/).

## Code availability

The custom pipelines used in this study are detailed in Methods, specific custom codes are available from the corresponding author upon request.

## Contributions

MJ and MPK conceived the research; MPK performed most of the experiments with help from CSC, LK, MK and RC. IA and GZ helped with the TMS design and data analysis. LK, MPK, EE, FR, MN, and TB helped with the data analysis. All authors have discussed the results and analyzed the data. LK, MPK and MJ wrote the manuscript with contributions from all authors. All authors read and approved the final manuscript.

## Ethics declarations

### Ethics approval and consent to participate

Not applicable.

### Consent for publication

Not applicable.

### Competing interests

The funders had no role in the study design, data collection and analysis, decision to publish, or preparation of the manuscript. The authors declare that they have no competing interests.

## Methods

### Construction of plasmids

The pdCas9-DNMT3A-Puro plasmid (Addgene #71667)^3^ was used as the backbone for all the epimodifier modules except CRISPRoff that we used the original designed cassette. To make a double reporter marker on the same backbone **(Suppl. Fig. 13)**, the T2A and puromycin modules were amplified from the pdCas9-DNMT3A-Puro by the primers containing the HindIII restriction sites. The amplified region was cloned on the pEGFP-C1 plasmid (gift from Alexander Espinosa). The whole cassette including the CMV enhancer, CMV promoter, EGFP, T2A and Puro was amplified by the primers containing the NotI restriction sites. Finally, after removing the Puro from the pdCas9-DNMT3A-Puro plasmid by EcoRI digestion and re-ligation, the CMV-GFP-T2A-Puro cassette was cloned on the pdCas9-DNMT3A plasmid (M3-dCas9-3A, Addgene #220237). The same procedure was applied to the control tool, consisting of dCas9 fused with the inactive DNMT3A (M3-dCAS9-DNMT3A, Addgene #220237). M3-dCAS9-DNMT3A-DNMT3A (dCas9-3A-3A, Addgene #218776) plasmid made by the amplification of the human DNMT3A catalytic domain from the pdCas9-DNMT3A-EGFP (Addgene #71666) plasmid using the primers containing the FseI restriction sites. For M3-dCAS9-DNMT3A-DNMT3L (dCas9-3A-3L, Addgene #218777), the murine DNMT3L catalytic domain and 27bp linker included FseI restriction site were amplified from pET28-Dnmt3a3L-sc27 plasmid (Addgene #71827). For the M3-dCAS9-DNMT3A-KRAB (dCas9-3A-KRAB, Addgene #218781), Krüppel-associated box (KRAB) transcriptional repression domain was amplified from the lenti-EF1a-dCas9-KRAB-Puro plasmid (Addgene #99372) with the primers containing the FseI restriction sites.

For M3-dCAS9-M.SssI (dCas9-M.SssI, Addgene #218782), the mutant version of prokaryotic methyl transferases M.SssI called MQ1^Q147L^ was amplified from pcDNA3.1-dCas9-MQ1(Q147L)-EGFP plasmid (Addgene #89634) by the primers containing the 27bp linker and SfiI and FseI restriction sites. For the M3-dCAS9-SunTag (dCas9-SunTag, Addgene #218783), 10XGCN4 region of the plasmid LLP457 pGK-dCas9-SunTag-BFP (Addgene #100957) was amplified by the primers containing SV40 NLS and PspxI and FseI restriction sites. Moreover, LLP252 pEF1a-NLS-scFvGCN4-DNMT3a plasmid (Addgene #100941) was used for the co-transfection with the generated dCas9-SunTag plasmid. R882H mutation has been conducted using site-directed mutagenesis (SDM) on dCas9-DNMT3A plasmid resulted in M3-dCAS9-mutDNMT3A(R882H) (dCas9-mut3A, Addgene #218788) cassette. Moreover, cassettes with UBC and EF1a promoters made by amplification of the relevant sequences and then cloning in dCas9-DNMT3A plasmid using the HiFi DNA Assembly Cloning Kit (NEB). Whole plasmid Sanger sequencing was performed to assess the cloning results (Eurofins Genomics, Germany).

### gRNA design and cloning

To have comparable results, *BACH2*, CXCR4 and non-targeting tested gRNAs were chosen from the previously published papers. For designing the targeted methylation sequencing panel, the off-target activity scores were checked by the CRISPOR online tool (http://crispor.tefor.net/crispor.py). To clone the gRNAs, we need the double stranded oligos with the appropriate overhangs for the BpiI sites. We added the cacc and aaac nucleotides to the 3’ of the sense and anti-sense gRNA oligoes, respectively. Annealing was performed using 12,5 ul (100 uM) sense oligo, 12,5 ul (100um) antisense oligo, 5ul fast digest buffer (Thermo Fisher Scientific) and H2O up to 50 ul. The mix is incubated by the following program: 95 °C for 5 min, gradually cool down (1 °C/min ramp) from 94 °C to 4 °C and storage in 4°C. All the gRNAs were cloned in the vectors in a single reaction including: 2 ul 10X fast digest buffer, 2 ul (10mM ATP), 500 ng plasmid, 1ul fast digest BpiI restriction enzyme (Thermo Fisher Scientific), 1,5 ul T4 DNA ligase (Thermo Fisher Scientific), 1ul (10 uM) annealed oligo and H2O up to 20 ul. The mix was incubated at 37 °C for 2 hours and inactivation was performed at 80 °C for 5-10 min. Bacterial transfection was performed by the TOP10 Electrocompetent E. coli cells. After colony selection, overnight culture and extraction of the plasmids, whole plasmid Sanger sequencing was performed to assess the cloning results (Eurofins Genomics, Germany). All the gRNA sequences are listed in the Supple. Table 1.

### In silico off-target prediction

In silico off-target prediction was performed utilizing a set of web-based tools (CRISPOR^47^ http://crispor.gi.ucsc.edu/, IDTDNA https://eu.idtdna.com/, Cas-OFFinder^48^ http://www.rgenome.net/cas-offinder/, CCTOP^49^ https://cctop.cos.uni-heidelberg.de) for both g8 and NTC gRNAs. Due to discrepancies in predicted off-targets arising from variations in tool functionalities and criteria, we compiled a comprehensive catalogue of these tools. This compilation facilitated the evaluation of predicted off-target occurrences within our RNA-seq and EPIC methylome datasets. While the allowable number of mismatches varied among these tools, we endeavored to extend the threshold to accommodate up to 5 mismatches whenever feasible.

### Transcription Factor Binding Site Analysis

To identify potential transcription factor binding sites (TFBS) within the genomic region targeted in the BACH2 TMS analysis, we used the JASPAR2022 database and Bioconductor packages in R. Transcription factor binding motifs were obtained from the JASPAR2022 CORE collection for human, and motif scanning was performed by aligning these motifs to the extracted sequences from the human genome (hg38). The presence and distribution of identified TFBS were systematically evaluated across the defined regions.

### Cell culture, transfection, and sorting

Human embryonic kidney HEK293T cells were maintained in Dulbecco’s Modified Eagle Medium (Sigma Aldrich, St. Louis, MO, USA) supplemented with 10% fetal bovine serum (Sigma Aldrich), 4 mM L-glutamine (Sigma Aldrich), 100 U/ml penicillin and 100 µg/ml streptomycin (Sigma Aldrich). Cells were incubated at 37^◦^C in a humidified 5% CO2-containing atmosphere. HEK293T cells were transfected using Lipofectamine 3000 Reagent (Invitrogen, Carlsbad, CA, USA) according to the manufacturer’s protocol. Briefly, cells were seeded in 6 well plates and were transfected the day after at around 80% confluency. The transfection cocktail was prepared as follows: 2-4µg of the plasmid DNA, 5 µl Lipofectamine™ 3000 Reagent, 2 µL/µg DNA of P3000™ Reagent and Opti-MEM™ Medium (Invitrogen). We used pKLV2-U6gRNA3(BbsI)-PGKpuro2ABFP (Addgene #67990) and replaced the BFP with mCherry (named as P3-pKLV2-U6gRNA(BbsI)-PGKpuro2A-mCherry) to express the gRNA when using the CRISPRoff cassette. For the SunTag construct, we added 400 ng αGCN4-DNMT3A vector to the cocktail. To ensure consistent level of transfection efficiency, cells were screened after 24 h for expression of the EGFP by fluorescent microscopy. To enrich the EGFP+ cells, FACS sorting was performed after 72 h and the sorted cells were subjected directly for DNA, RNA and protein extraction using AllPrep DNA/RNA/Protein Mini Kit (QIAGEN, Venlo, Netherlands).

### DNA methylation analysis

#### Pyrosequencing

DNA treatment with sodium-bisulfite that leads to the conversion of all unmodified cytosine to uracil while 5mC are resistant to deamination induced by bisulfite, and are preserved during the PCR amplification by primers designed on the converted DNA ^50^. 200 ng of extracted DNAs were subjected to bisulfate conversion using EZ DNA Methylation-Gold Kit (Zymo Research Europe, Freiburg, Germany) according to the manufacturer’s protocol. The bisulfite converted DNA (∼10 ng) was amplified by PyroMark PCR kit (QIAGEN, Hilden, Germany) according to the manufacturer’s instructions. The PCR was performed using 5 pmol of forward and 5′-biotinylated reverse primers with the following cycles: initial denaturation for 15 min at 95◦C; 50 cycles of 30 s at 94◦C, 30 s at 48◦C (*BACH2*) or 50◦C (*CXCR4*) or 60 (*IL6ST*) and 30 s at 72◦C; final extension for 10 min at 72◦C. Finally, PCR products were sequenced by PyroMark Q24 Advanced pyrosequencing system (QIAGEN) using PyroMark Gold 96 reagent kit (QIAGEN). All pyrosequencing PCR primers and probes were designed by PyroMark Design software (QIAGEN) and are listed in the Supple. Table 1.

#### Targeted methylation sequencing (TMS)

For high-resolution analysis of the methylation status of all CpG sites surrounding the target gRNAs and their potential off-target loci predicted by CRISPOR, we employed QIAseq Targeted Methyl Panel technology. The primer panels to enrich these regions were designed by QIAGEN (Hilden, Germany) both for the *BACH2* (258 CpGs, 33 loci, 22,768 bp) and *CXCR4* (212 CpGs, 13 loci, 10,871 bp) gRNAs (Supple. Table 2). Briefly, 110 ng DNA was bisulfite converted using EpiTect Fast Bisulfite Conversion Kit according to the manufacturer’s instructions. Converted DNA fragments were attached to unique molecular identifier (UMI) and libraries were created using the QIAseq Targeted Methyl Panel protocol and were amplified for 18 cycles. Library quality was assessed using an Agilent 2100 Bioanalyzer (Agilent Technologies, Inc.) and quantified using KAPA Biosystem’s next-generation sequencing library qPCR kit. Libraries were pooled in equimolar concentrations and 10 pM pooled library was sequenced using NovaSeq 6000 as paired end 150 sequencing (Illumina, San Diego, CA, USA). Data were analyzed using the QIAGEN CLC Genomics Workbench (QIAGEN, Aarhus, Denmark) and if the coverage of the context was less than 10X, we assigned methylation values to "missing".

#### EPIC

The Illumina EPIC array was used by NGI in Uppsala, Sweden to measure DNA methylation in all samples collected on day 3 p.t, as well as in a subset of selected samples with superior DNA methylation deposition performance from days 7 and 30 p.t, along with appropriate controls. Data analysis was performed using the Illumina default procedure implemented in the Bioconductor minfi package ^51^. We have defined a differentially methylated region (DMR) as a consecutive sequence of two CpGs exhibiting a methylation increase of over 10% when they are in a distance less than 2 kb.

### Gene expression analysis

#### Real-time PCR

Genomic DNA, RNA and protein were extracted simultaneously from the fresh EGFP sorted cells using AllPrep DNA/RNA/Protein Mini Kit (QIAGEN). Reverse transcription was performed using 100 ng of the total RNA by iScript cDNA synthesis Kit (Bio-Rad) by random hexamer and poly A primers. Real-time PCR was performed on a BioRad CFX384/C1000 Real-Time Detection System using Cyber Green (Bio-Rad) with the following cycling protocol: 95 °C:3 min, followed by 40 cycles of 95 °C:10 s, 60 °C: 30 s and 72 °C:30 s. All primers are listed in Supple. Table 1. The relative expression of the genes were normalized to *HPRT* as the housekeeping endogenous control. Relative quantification of mRNA levels was performed using the ΔΔCt method as described by Livak. Statistical significance was determined using a two-tailed t-test and a p-value <0,05 was considered significant. All statistical analysis was performed by GraphPad Prism 6.01 (GraphPad Software, Inc., San Diego, CA, USA).

#### RNA-sequencing

The RNA-seq libraries for day 3 p.t. were constructed according to the manufacturer’s protocol using the Illumina® Stranded Total RNA Prep, Ligation with Ribo-Zero Plus (Illumina, CA, USA). The quantity and quality of the libraries were assessed using a Qubit fluorimeter (Thermo Fisher Scientific) and Bioanalyzer DNA Electropherogram (Agilent Technologies). Sequencing was carried out on Illumina NovaSeq 6000 with 150-base pair paired end reads (NGI, Sweden). The RAW RNA sequence files were used to extract the fastq data, which underwent quality control checks using the multiqc software to prepare them for alignment ^52^. Subsequently, the resulting fastqc files were aligned and annotated using the STAR aligner and Stringtie software ^53^, incorporating human hg38 refseq information from UCSC. Finally, the analysis of the count matrix was carried out using bash and Python and reads were mapped to the human reference genome GRCh37 (hg19) assembly. To filter out low-expressed transcripts, we have considered 25% of the samples have at least five count per transcript. The counts were normalized by CPM using the TMM method and log2 fold change (logFC) was calculated based on the normalized CPM counts. We chose logFC ≥ 1 or logFC < −1 and adjusted *p* value < 0.05 as the screening criteria. mRNA-seq analysis was conducted to investigate transcriptome changes associated with methylation alterations in samples collected at days 3, 7, and 30 p.t. (Novogene). All transcriptome data were deposited in the SRA NCBI data repository (Bioproject: XXX).

#### Western blotting

Adding the epimodifiers with different sizes to the dCas9 protein making fusion proteins with a range of seizes. To check the expression and the size of the fusion dCas9-Epimodifiers, western blotting using an antibody against Cas9, was performed. In both experiments, CelLytic M (Sigma) buffer supplemented with Halt Protease and Phosphatase Inhibitor Cocktail (Thermo Fisher Scientific) was used for cell lysis. The protein concentration was measured using Pierce™ 660 protein assay reagent (Thermo Fisher Scientific #22660). The supernatant was mixed with Laemmli sample buffer 4X (Bio-Rad, #1610747) supplemented with 50mM DTT (Sigma-Aldrich, #10197777001) and heated at 95°C for 5min. Equal quantities of proteins from each sample (20 μg) were separated on mini-Protean® TGX™ mini pre-cast gels (Bio-Rad, #4561023) at 120V. The proteins were transferred to a PVDF transfer membrane (0.45 μm, ThermoScientific #88518) by using a Trans-Blot®Turbo™ transfer system (Bio-Rad) at a constant current of 2.5A for 30 minutes. The membranes were blocked using TBS/0.1% Tween (Sigma-Aldrich) with 5% nonfat dried milk (Panreac AppliChem). The membrane was incubated overnight at 4°C with the primary antibody (mouse anti-CRISPR-spCas9 antibody 1:1000, ab191468, Abcam, Cambridge, UK), washed in TBS/0.1% Tween and followed by an one hour incubation with a secondary antibody conjugated to HRP (goat Anti-Mouse IgG H&L (HRP) 1:3000, #7076, Cell Signaling). Beta-actin was used as a loading control and the antibody was directly conjugated to HRP (Anti-Beta-Actin conjugated with HRP 1:25,000, Sigma-Aldrich, A3854 (clone AC-15). The band detection was achieved by using Clarity™ Western ECL substrate (Bio-Rad). The band intensities were analyzed with Image Lab™ software (Bio-Rad).

## Notes

### Competing Interest Statement

The authors have declared no competing interest.

