## Supplementary Figures for "Systematic comparison of dCas9-based DNA methylation epimodifiers over time indicates efficient on-target and widespread off-target effects"

#### Supplementary Figure 1

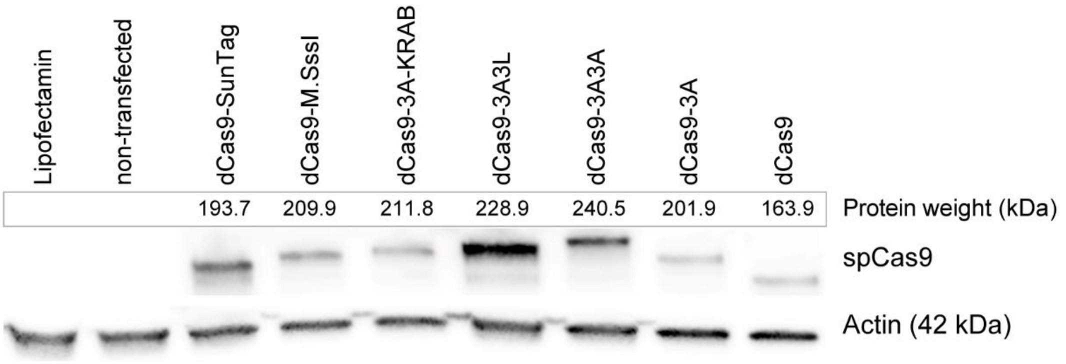


**Supplementary Fig 1.** Western blot of dCas9-epimodifiers.
The expression and size of the fusion between dCas9 and each epimodifier were assessed using the antibody against Cas9. An anti-actin antibody served as a loading control.

#### Supplementary Figure 2

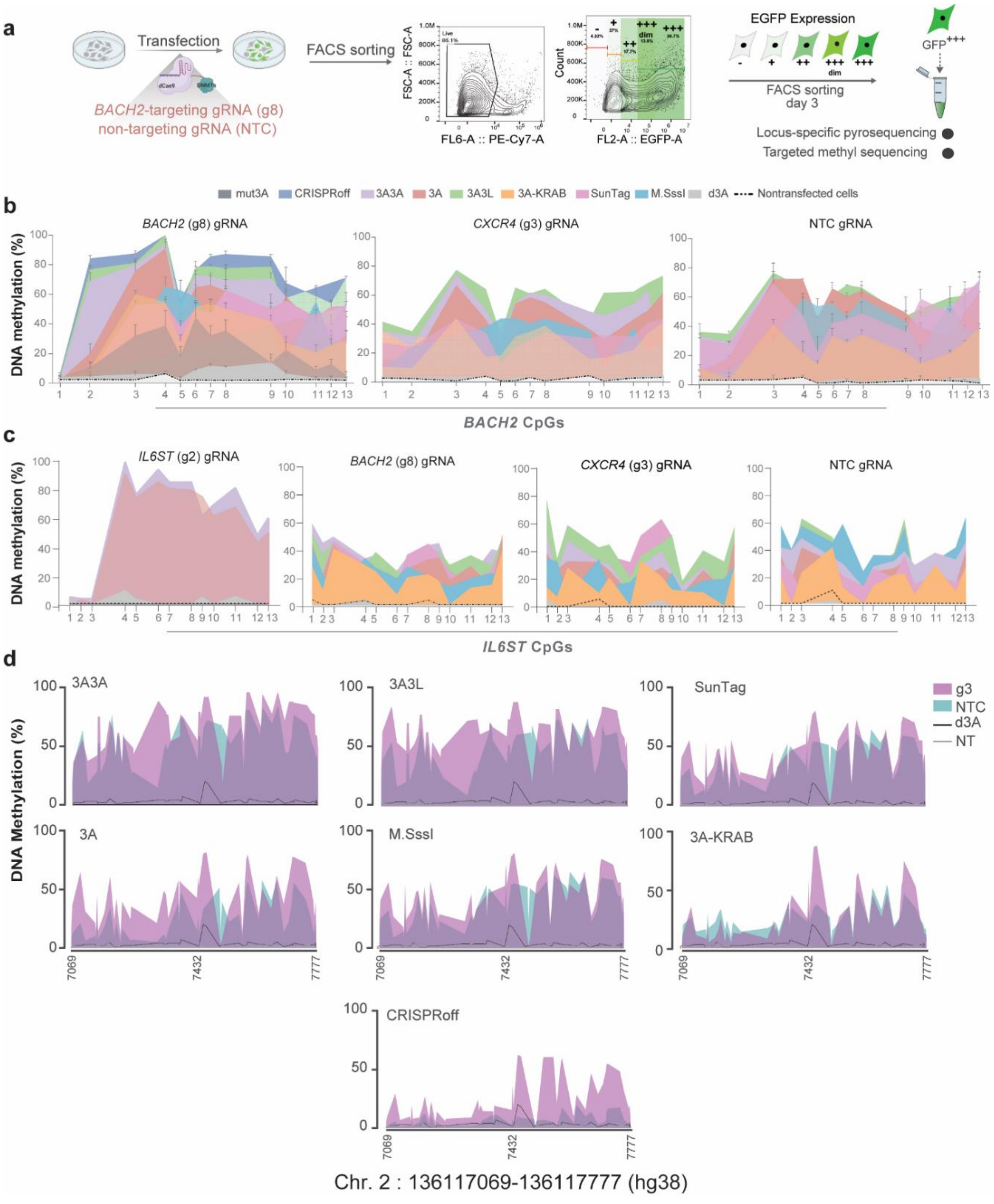


**Supplementary Fig. 2.** Early targeted methylation deposition by dCas9-epimodifiers.
**a.** Schematic figure showing details of the experimental design and sorting strategy. We selected cells with the highest GFP protein levels, which correlates with the highest expression of dCas9-epimodifier constructs. **b.** As evidenced by targeted methyl sequencing (TMS, Fig. 1), our pyrosequencing data validate methylation deposition at the targeted site (*BACH2* promoter). By utilizing irrelevant gRNAs (*CXCR4*-g3), as well as the non-targeting control gRNA (NTC), we observed significant methylation deposition in the *BACH2* promoter, indicating a high level of specific gRNA-independent off-target methylation. **c.** Similar on- and off-target effects were observed when analyzing additional loci, i.e., *IL6ST*, using the targeted gRNA (*IL6ST*-g2), as well as the irrelevant gRNAs (*BACH2*-g8, *CXCR4*-g3) and NTC. **d.** DNA methylation of CpGs at *CXCR4* promoter assayed using pyrosequencing 3 days post-transfection with the epimodifiers expressing either *CXCR4*-targeting gRNA3 (g3, purple color) or non-targeting gRNA (NTC, light green color). Methylation levels in the control cells including cells dCas9-deactivated DNMT3A (d3A)- or non-transfected cells (NT) are depicted by the black and grey lines, respectively.

#### Supplementary Figure 3

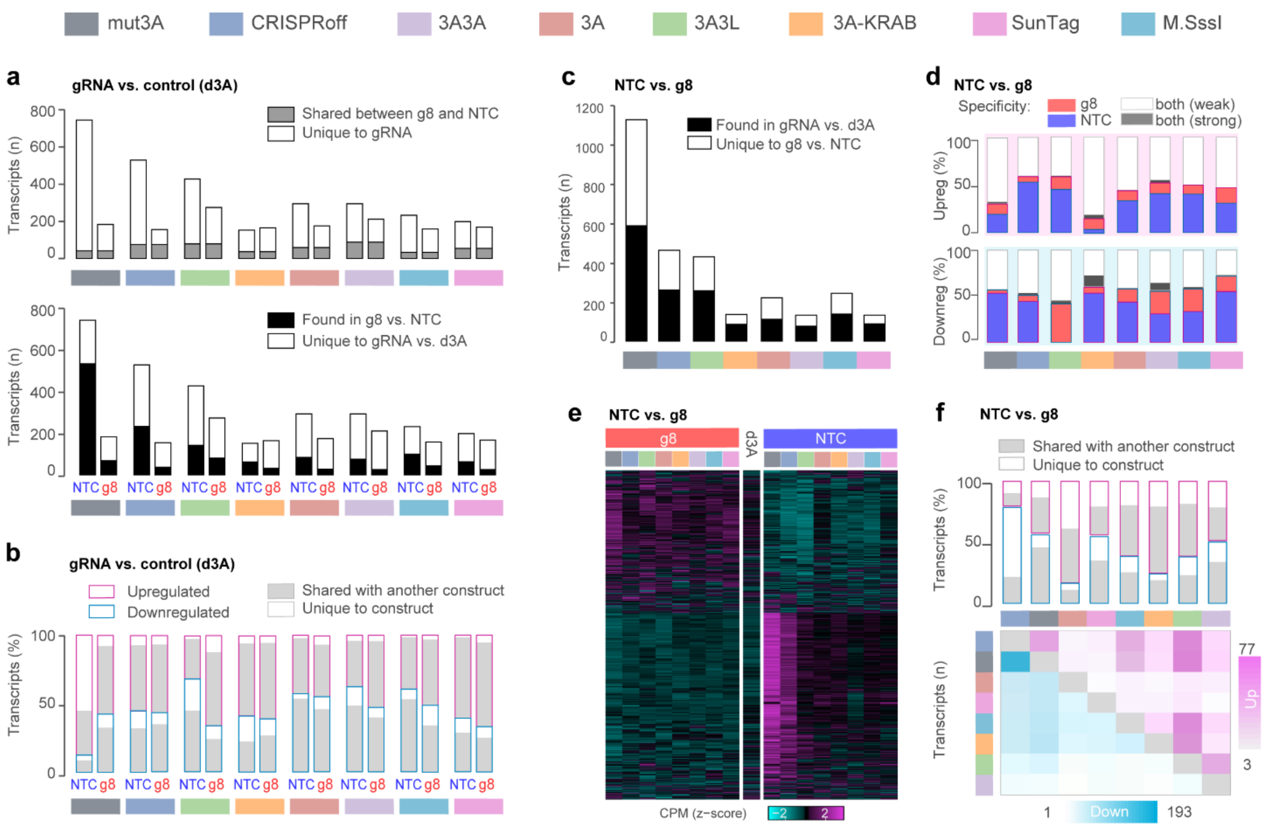


**Supplementary Fig. 3.** Early transcriptional effects of dCas9-epimodifiers.
**a**. Number of transcripts differing between either non-targeting gRNA control (NTC) or *BACH2*-targeting gRNA (g8) and the control dCas9-deactivated DNMT3A (d3A), with $|\log_2 FC| > 1$. The fraction of them that is shared between NTC and g8 in each construct is highlighted in grey color (upper graph) and the number of transcripts found also in the comparison g8 vs. NTC is shown in black (lower graph). **b**. Percentage of transcripts differing between NTC or g8 and the control d3A ($|\log FC| > 1$) that is upregulated (pink) or downregulated (blue). The fraction shared with at least one other epimodifier is depicted in grey color. **c**. Total number of the transcripts and the fraction of them (black color) that display difference ($|\log FC| > 1$) between either NTC or g8 and the control d3A. **d**. Origin of the upregulated (upper graph) and downregulated (lower graph) transcripts found in g8 vs. NTC ($|\log FC| > 1$). The fraction of changes arising from difference between one gRNA, i.e. NTC or g8, and the control d3a solely is represented in blue and red colors, respectively. The fraction of transcripts with $|\log(g8/NTC)| > 1$ that show weak ($|\log FC| < 1$) or strong ($|\log FC| > 1$) differences between both NTC and g8 and the control d3A. **e**. Heatmap of the counts (CPM, count per million) of the transcripts differing between g8 and NTC with $|\log_2 FC| > 1$. **f**. Proportion (%) of upregulated (left) and downregulated (right) transcripts with absolute logFC (g8/NTC) > 1 that are shared with at least one other epimodifier and pairwise number of shared upregulated (pink color) and downregulated (blue color) transcripts.

#### Supplementary Figure 4

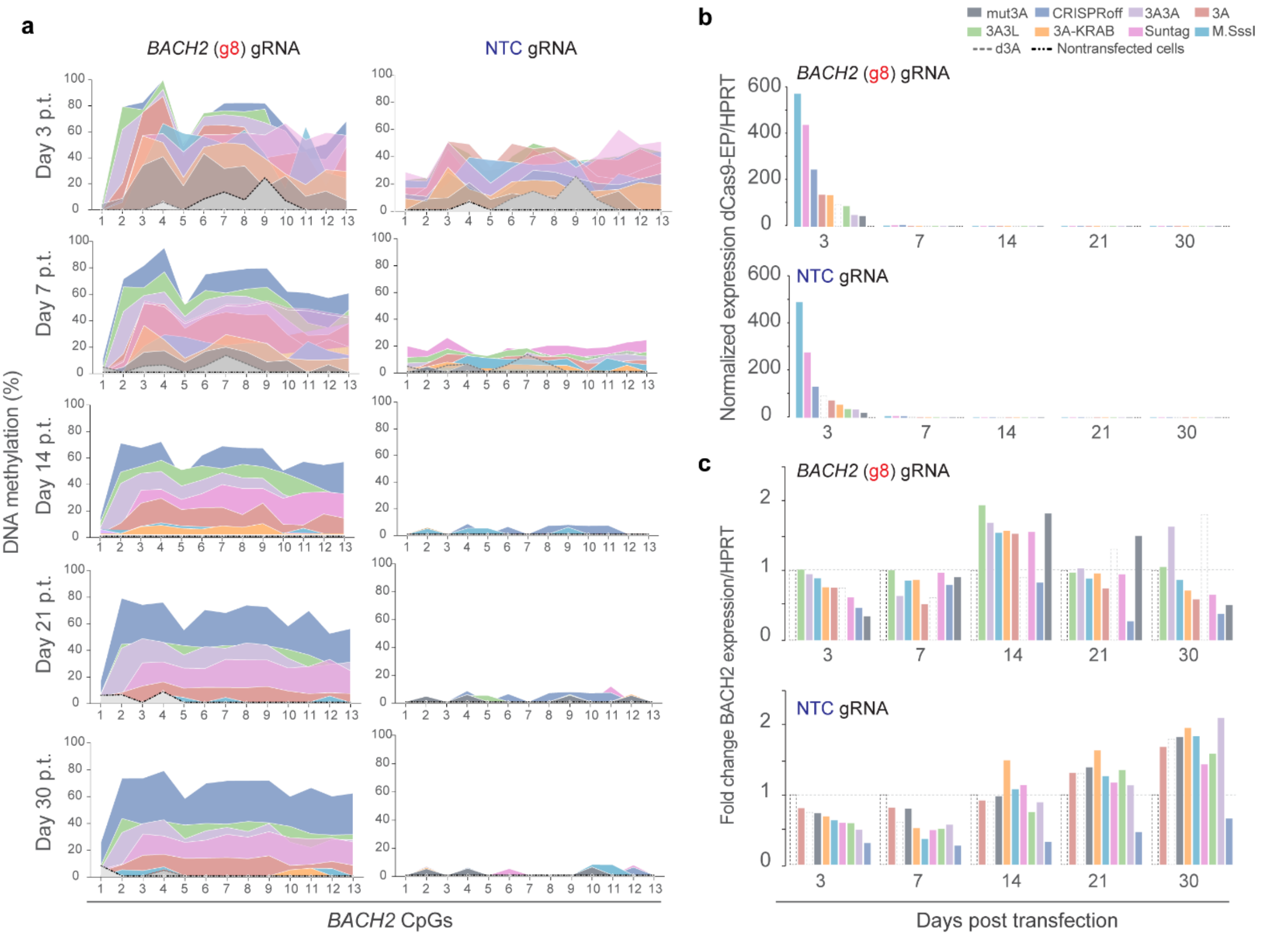


**Supplementary Fig. 4.** Evaluation of the stability of the deposited methylation over time.
**a.** Pyrosequencing data from the *BACH2* locus show high methylation deposition at day 3 post-transfection (p.t.) both for *BACH2*-targeting gRNA (g8) and non-targeting gRNA control (NTC) for all dCas9-epimodifiers. While CRISPRoff, dCas9-3A3L, and dCas9-3A3A are the most efficient constructs for stable methyl-deposition, the residual effect of NTC persisted until day 7 p.t. and subsequently decreased to control levels for all constructs. **b.** Normalized expression levels of all epimodifiers were tracked over time. Aligned with the nature of the transient transfection, no detectable expression of the dCas9-epimodifiers was observed from day 7 p.t. in all conditions. **c**. Normalized expression levels of the *BACH2* gene, over time. The most potent methylating constructs, CRISPRoff, dCas9-3A3L and dCas9-3A3A, also exhibited the strong *BACH2* repressing capacity as revealed by RNA-seq data (Fig. 3f).

#### Supplementary Figure 5

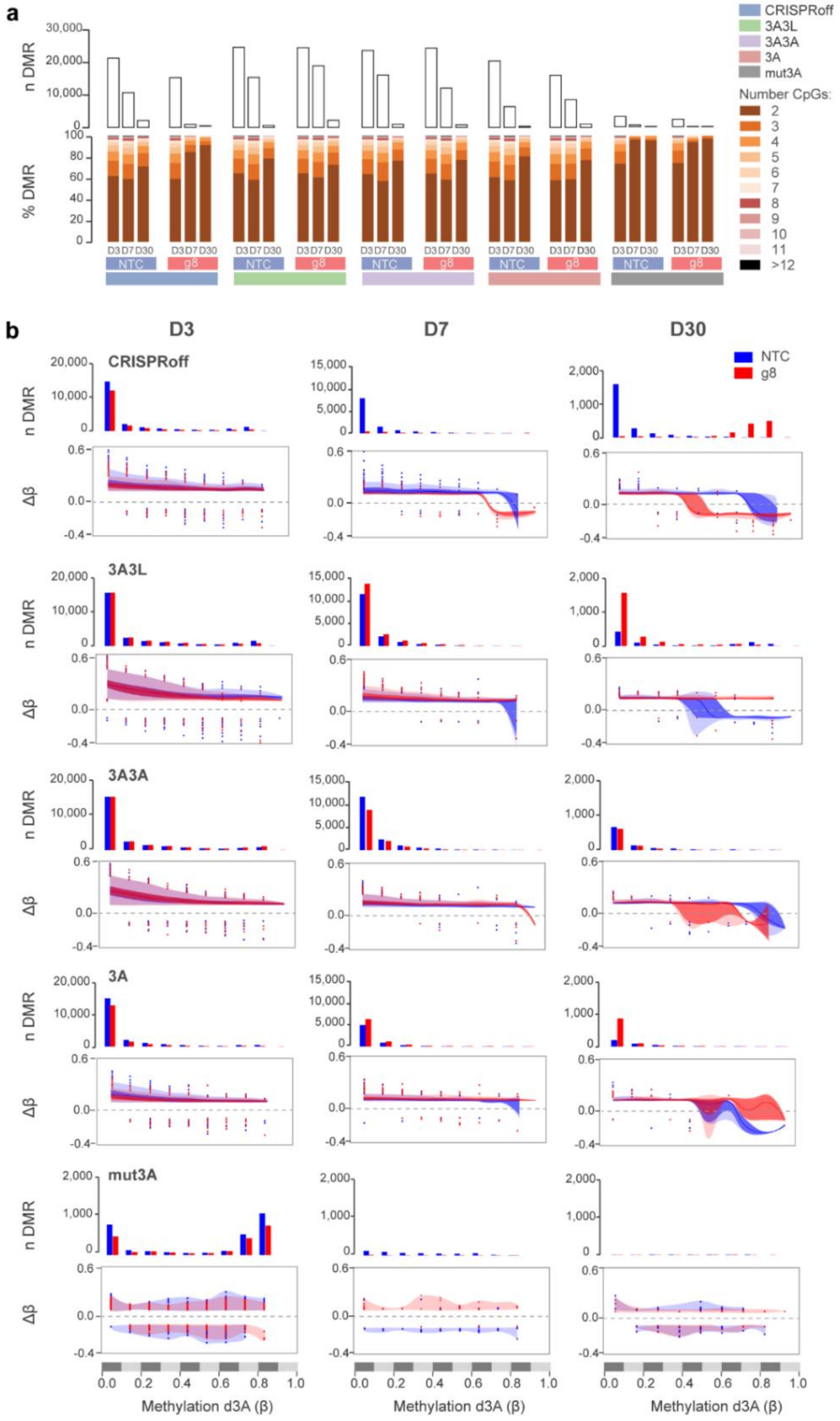


**Supplementary Fig. 5.** Characteristics of off-target methylation deposition.
**a**. Number of total differentially methylated regions (DMRs, |Δβ| > 0.1) between either the non-targeting control gRNA (NTC) or *BACH2*-targeting gRNA (g8) and dCas9-deactivated DNMT3A (d3A) control at days 3 (D3), 7 (D7) or 30 (D30) post-transfection (p.t.). The proportions of DMRs according to the number of CpGs included are depicted in different orange and red color gradients. **b**. The number of DMRs and average methylation differences in relation to the basal methylation level (0.2 β-value bins) in the control deactivated DNMT3A (d3A) construct are shown for each dCas9-epimodifier and time point.

#### Supplementary Figure 6

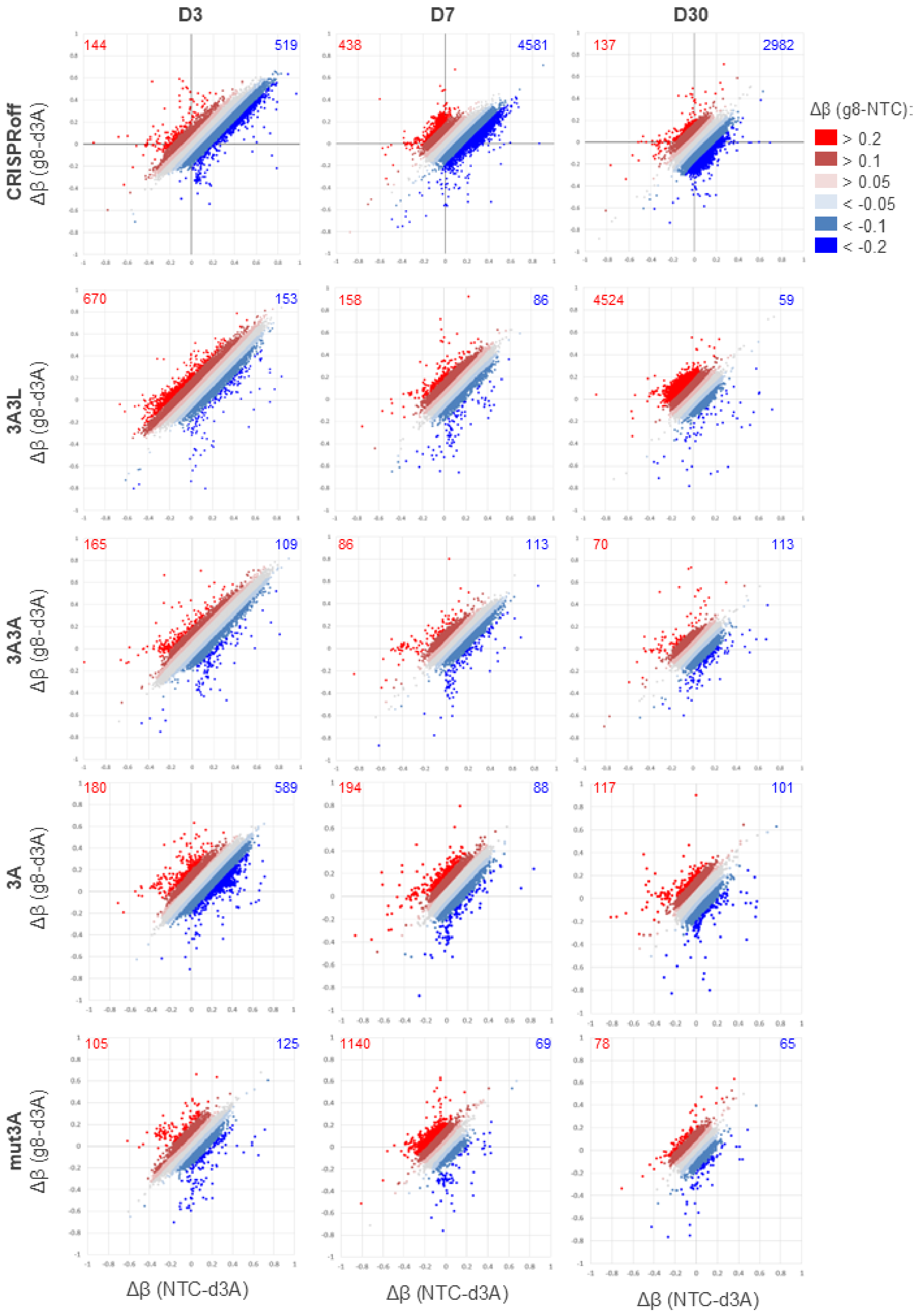


**Supplementary Fig. 6**. Differentially methylated CpG positions.
Correlation between the methylation changes induced by *BACH2*-targeting gRNA (g8, y-axis) or non-targeting control gRNA (NTC, x-axis) in comparison to the control dCas9-deactivated DNMT3A (d3A) control at days 3 (D3), 7 (D7) or 30 (D30) post-transfection. The hypermethylated and hypomethylated CpGs differing between g8 and NTC are highlighted in red and blue colors, the numbers refer to the number of CpGs displaying $|\Delta\beta_{\text{g8-NTC}}| > 0.1$.

#### Supplementary Figure 7

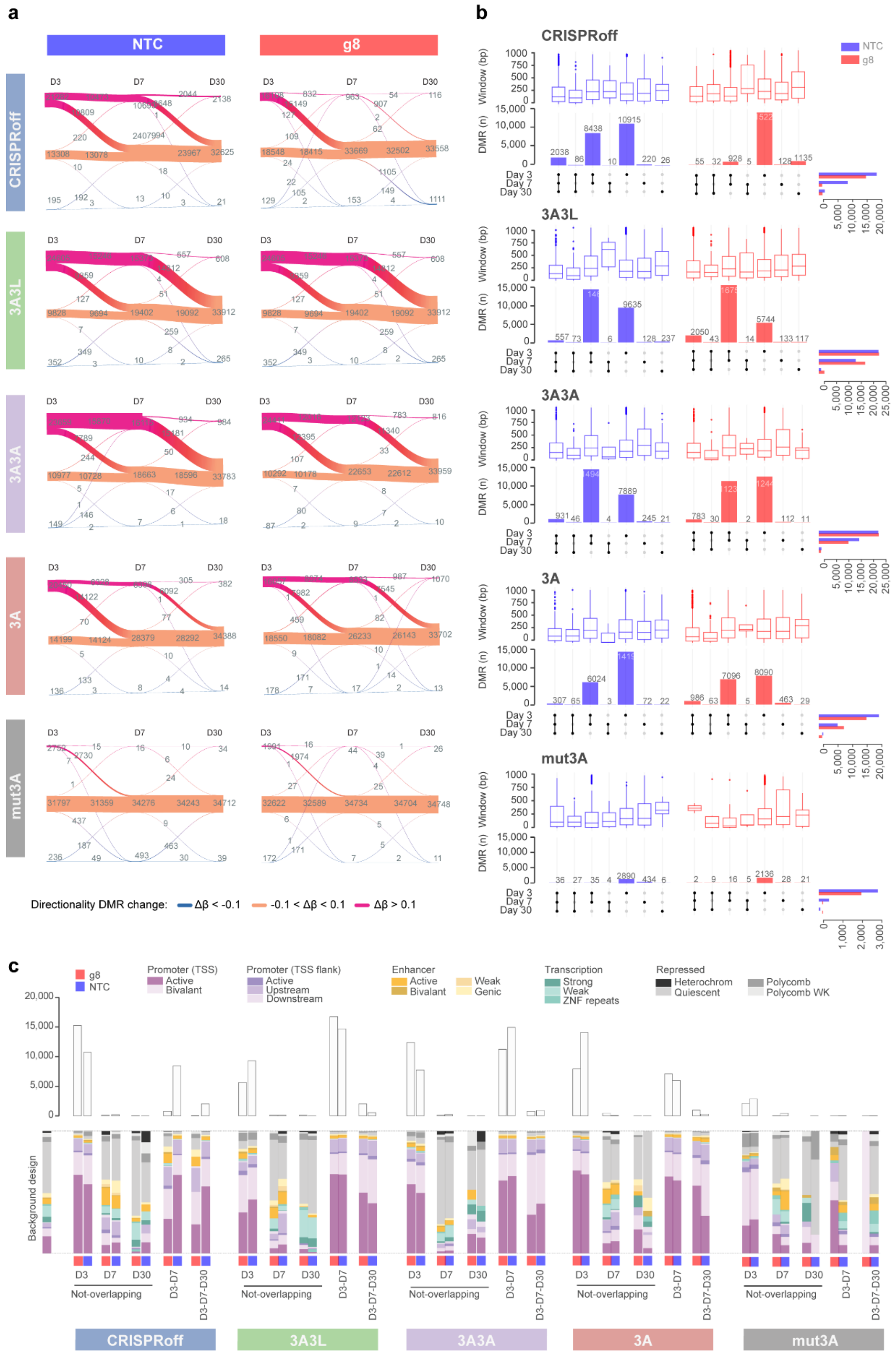

**Supplementary Fig. 7.** Characteristics and stability of off-target methylation deposition over time.
**a**. The number and size (window in bp) of differentially methylated regions (DMRs) overlapping between time points or specific for each time point are shown for each epimodifier and gRNA. **b**. Numbers of hypermethylated (Δβ > 0.1) and hypomethylated (Δβ < -0.1) DMRs over time, i.e., days 3 (D3), 7 (D7) or 30 (D30) post-transfection. **c**. Distribution of DMRs that are stable throughout time (overlapping between days 3 and 7 or days 3, 7 and 30) or DMRs that are specific for each time point across regulatory states in the HEK293T genome (chromatin HMM segmentation was obtained from the International Human Epigenome Consortium).

#### Supplementary Figure 8

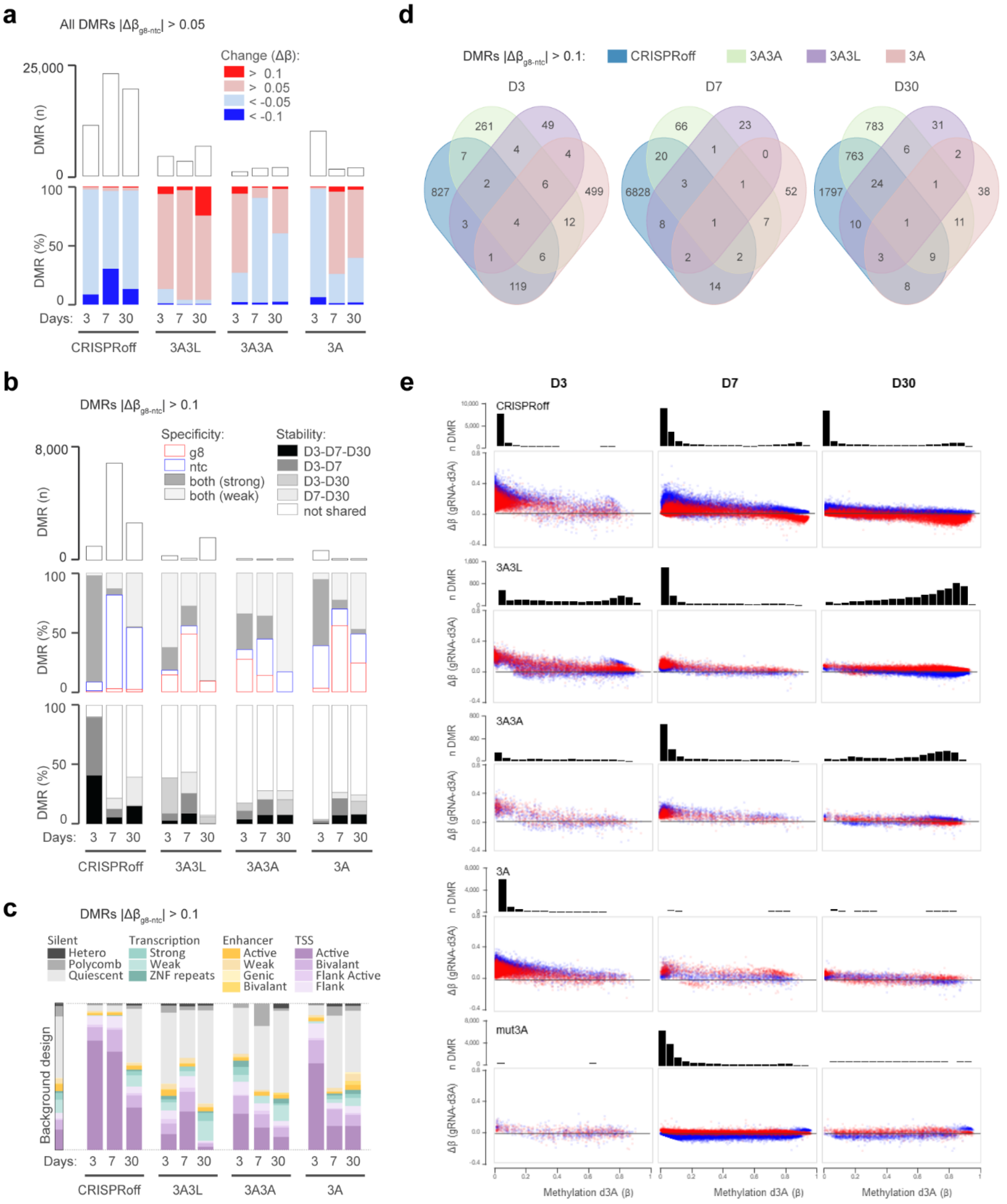


**Supplementary Fig. 8**. Characteristics of gRNA-specific off-target methylation deposition over time.
**a**. Total number (upper panel) and directionality of the changes (lower panel) at the differentially methylated regions (DMRs) that differ between *BACH2*-targeting gRNA (g8) or non-targeting control gRNA (NTC) in comparison to the control dCas9-deactivated DNMT3A (d3A) control at days 3 (D3), 7 (D7) or 30 (D30) post-transfection. **b**. Total number of DMRs (upper panel), origin of the changes (middle panel) and stability (lower panel) of the most robust changes (|Δβ| > 0.1) between g8 and NTC. **c**. Distribution of DMRs across regulatory states in the HEK293T genome (chromatin HMM segmentation was obtained from the International Human Epigenome Consortium). **d**. Number of DMRs overlapping between dCas9-epimodifiers at each time point. **e**. Distribution of DMRs according to the basal level of methylation in the control deactivated DNMT3A (d3A) condition.

#### Supplementary Figure 9

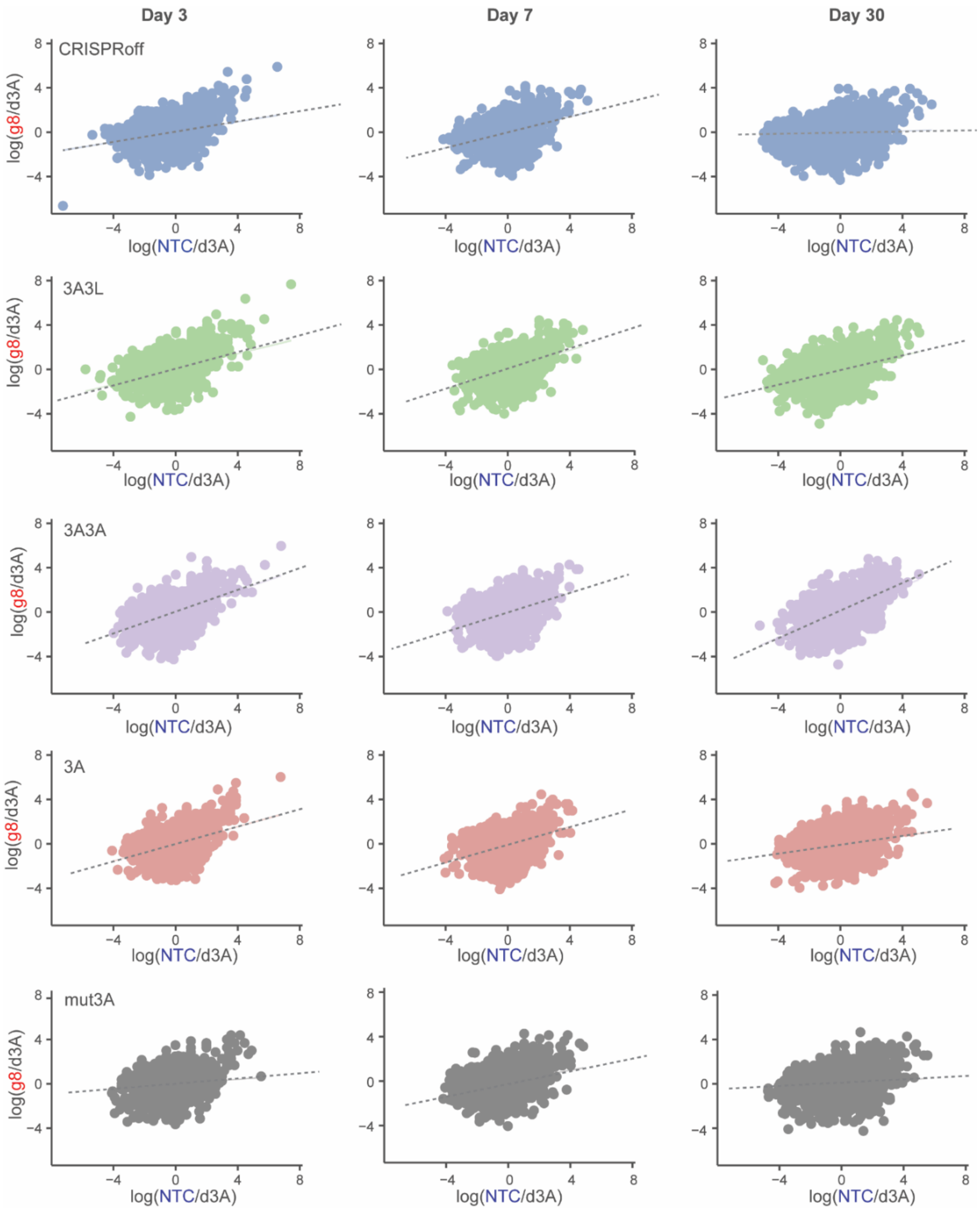


**Supplementary Fig. 9**. Transcriptional effects of the four potent dCas9-epimodifiers over time. The X and Y axes represent log2 gRNA/control (dCas9-deactivated DNMT3A, d3A) for *BACH2*-targeting gRNA (g8) and non-targeting control gRNA (NTC) 3, 7 and 30 days post-transfection, respectively.

#### Supplementary Figure 10

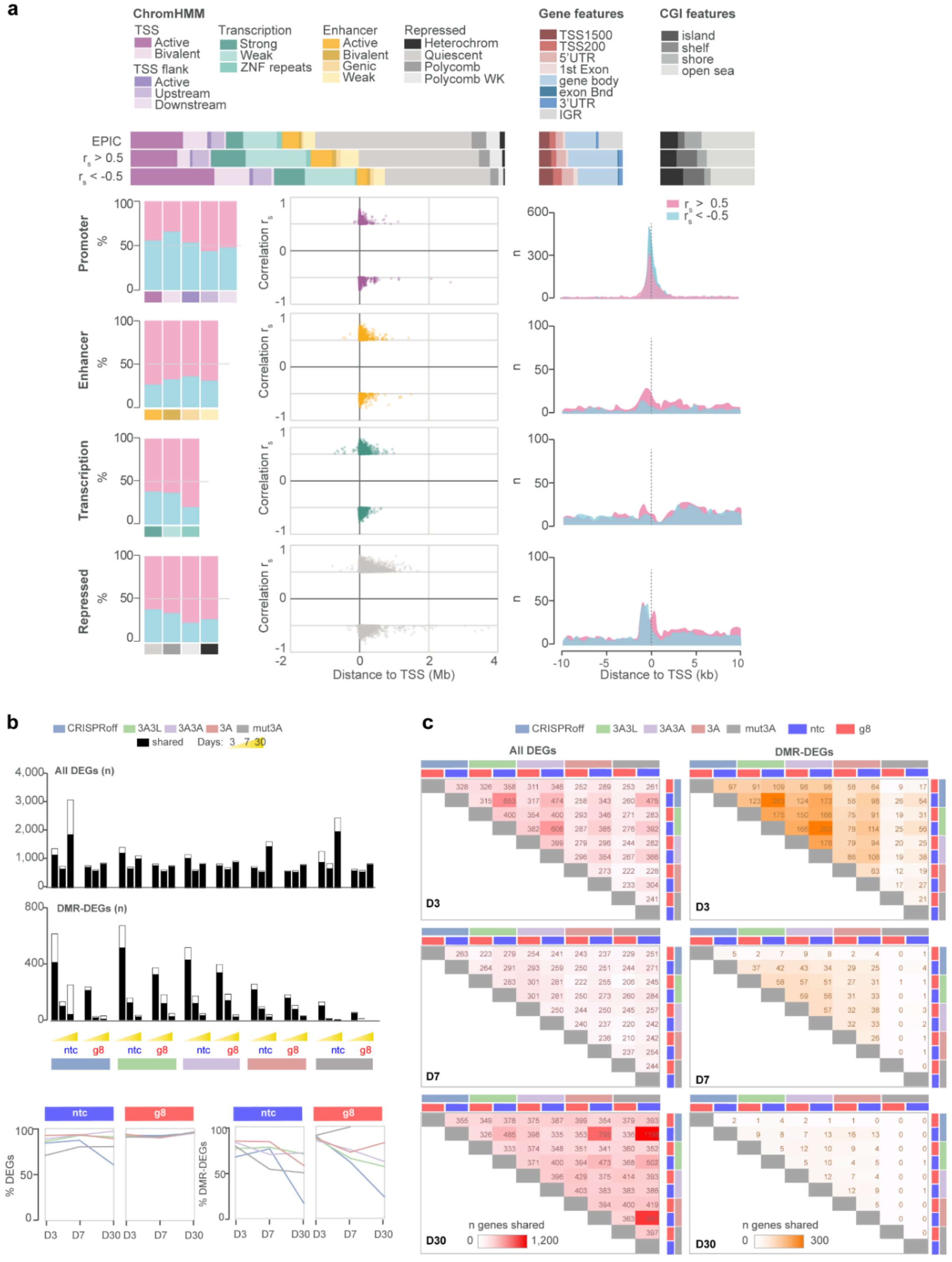


**Supplementary Fig. 10**. Colocalized methylation-expression changes over time.
**a.** Distribution of the 26,240 CpG-gene pairs displaying correlation (Spearman $P < 0.05$ and $|rho (r_s)| > 0.5$) across ChromHMM-, gene- and CpG island-related features and in relation to distance to transcription start site (TSS). Chromatin HMM segmentation was obtained from the International Human Epigenome Consortium. **b.** Number and percentage of all altered genes and differentially methylated regions that colocalized with differentially expressed genes (DMR-DEGs, $|\Delta\beta| > 0.1$ and $|\log_2 FC| > 1$) that are shared between at least two dCas9-epimodifiers. **c.** Details of the number of DEGs and DMR-DEGs shared between dCas9-epimodifiers 3 (D3), 7 (D7) and 30 (D30) days post-transfection.

### Supplementary Figure 11

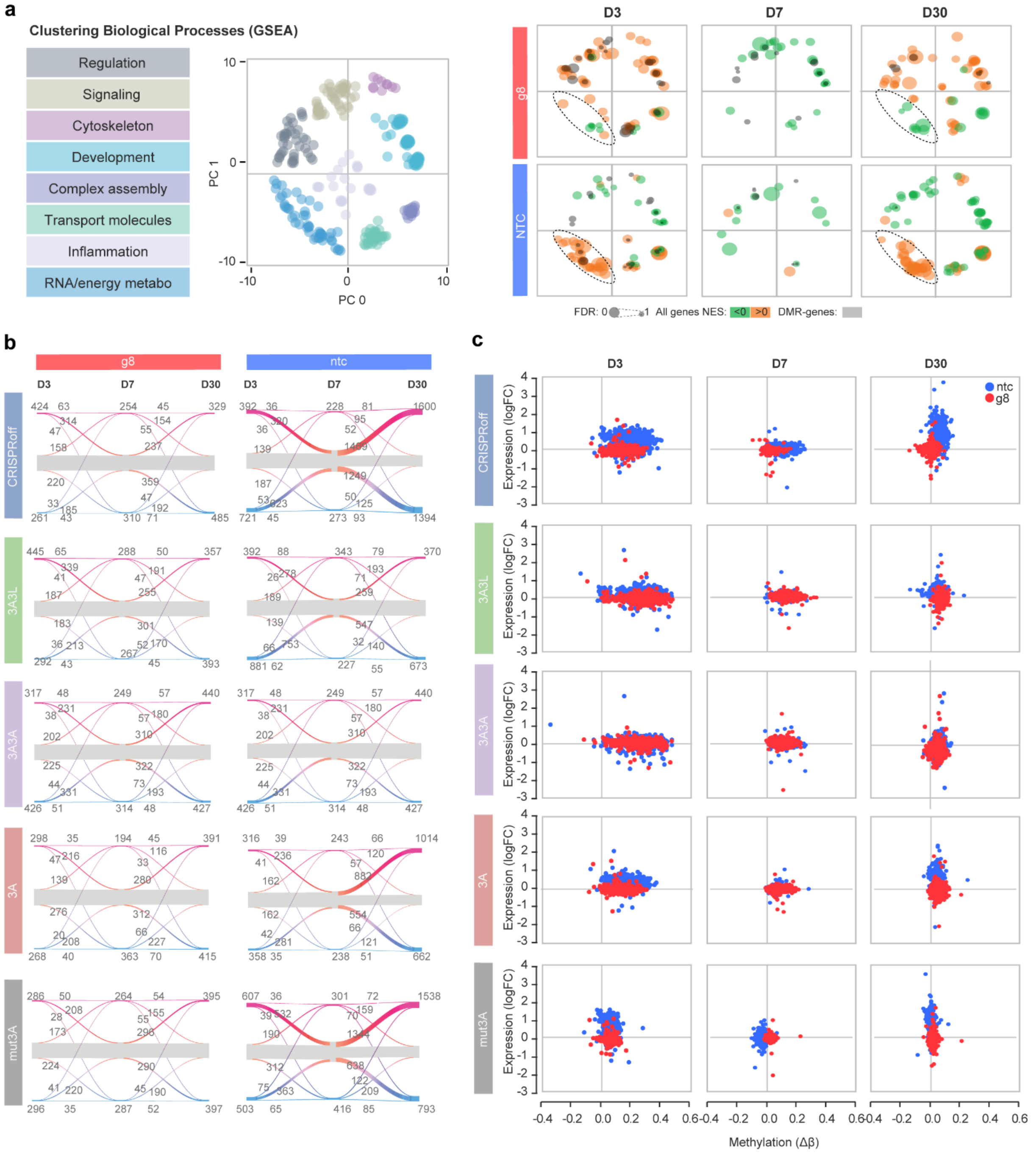


**Supplementary Fig. 11**. Stability of transcriptional changes over time.
**a.** Clustering of all ‘Biological processes’ terms (Gene Set Enrichment Analysis, GSEA, unadjusted P < E-03) from altered genes at days 3 (D3), 7 (D7) and 30 (D30) post-transfection (genes displaying $|log_2FC| > 1$) using multidimensional scaling according to semantic similarities and visualized using REVIGO. The circle size illustrates –log10 (FDR), green/orange colors represent negative and positive normalized enrichment score (NES), respectively. GSEA terms associated to differentially methylated regions (DMRs) that colocalized with differentially expressed genes (DMR-DEGs) are depicted in grey. The cluster differing the most between *BACH2*-targeting gRNA (g8) and non-targeting control gRNA (NTC) is highlighted by a dotted line. **b**. River plots of the number of genes varying across time points. **c**. Correlation between methylation changes (Δβ) and expression changes (logFC) between epimodifers and the control dCas9-dDNMT3A for 486 out of the 647 genes included in the top GSEA terms for which DMR info was available**.**

#### Supplementary Figure 12

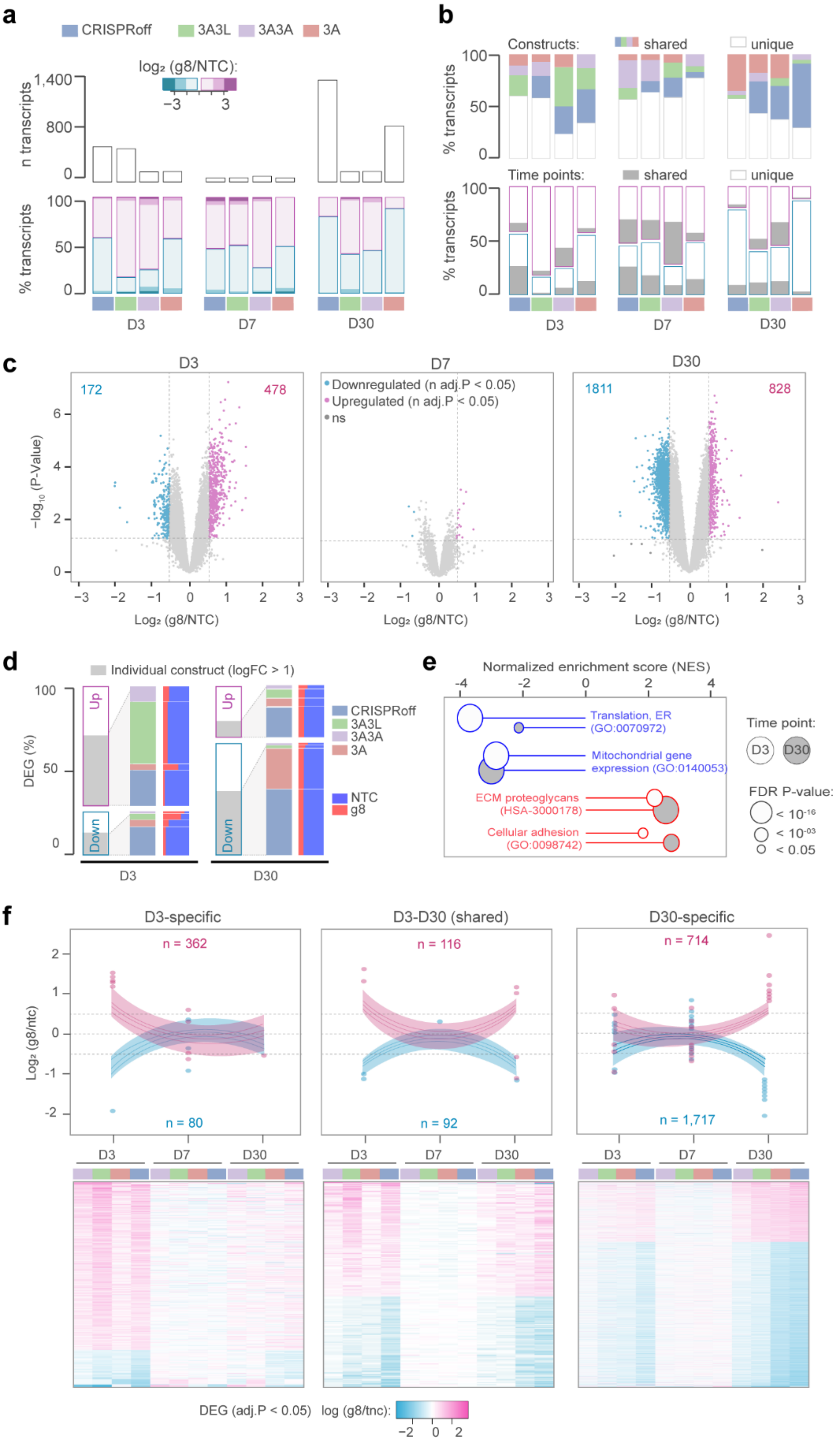

**Supplementary Fig. 12**. Characteristics of gRNA-specific transcriptional changes over time.
**a.** Total number of transcripts dysregulated between *BACH2*-targeting gRNA (g8) and non-targeting control gRNA (NTC) (absolute logFC > 1) and fraction of those that are upregulated (pink) and downregulated (blue). **b**. Percentage of differentially expressed transcripts between g8 and NTC that are shared with gRNA-epimodifier condition (upper graph) or that are overlapping between other time points (grey, lower graph). **c**. Volcano plot of the significantly upregulated and downregulated transcripts between g8 (n = 4 epimodifiers) and NTC (n = 4 epimodifiers). Nominally significant (P-value < 0.05) upregulated and downregulated transcripts ($|\log_2 FC| > 0.5$) depicted in purple and blue colors, respectively. The numbers refer to the significant differentially expressed genes (DEGs) after adjustment for multiple testing (adj. P < 0.05). **d**. Annotation of gRNA-specific DEGs (adj. P < 0.05) at days 3 (D3) and 30 (D30) post-transfection (p.t.) according to the directionality of change, the individual dCas9-epimodifier construct, gRNA and methylation changes. **e.** Most significantly enriched biological processes of the DEGs (adj. P < 0.05) at days 3 and 30 p.t.. Gene set enrichment analysis GSEA was used to calculate the normalized enrichment score (NES) and P-values. **f**. Expression changes (logFC) of the unique and shared upregulated (pink) and downregulated (blue) DEGs between g8 (n = 4 epimodifiers) and NTC (n = 4 epimodifiers) DEGs (adj. P < 0.05) found at day 3 and day 30 with corresponding heatmap depicting each dCas9-epimodifier. The graph lines depict the median +/- quartiles and the dots represent >1.5 interquartile range outliers.

#### Supplementary Figure 13

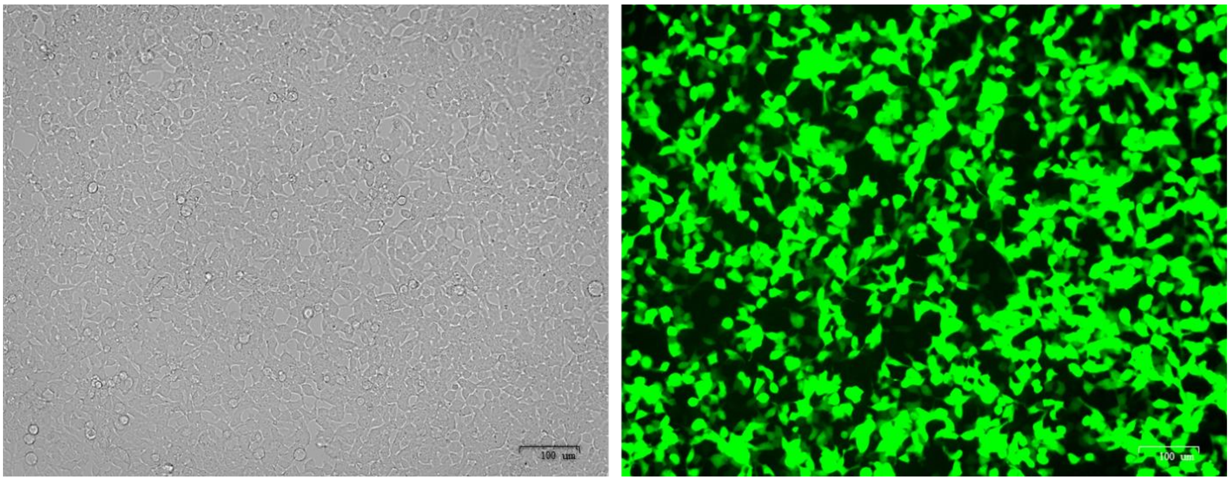


**Supplmentary Fig. 13.** Strategy to isolate transfected cells.
All dCas9-epimodifier constructs express both GFP and Puromycin under the control of a CMV promoter enabling enrichment and isolation of transfected cells only. Right, successfully transfected HEK293T cells with dCas9-DNMT3A expressing GFP, 3 days post-transfection; left, brightfield microscopy image of the same field showing all HEK293T cells in culture. Scale bar: 100 µm.
